# Sites of Circadian Clock Neuron Plasticity Mediate Sensory Integration and Entrainment

**DOI:** 10.1101/794537

**Authors:** MP Fernández, HL Pettibone, CJ Roell, CE Davey, KV Huynh, SM Lennox, BS Kostadinov, OT Shafer

## Abstract

Networks of circadian timekeeping in the brain display marked daily changes in neuronal morphology. In *Drosophila melanogaster*, the striking daily structural remodeling of the dorsal medial termini of the small ventral lateral neurons has long been hypothesized to mediate endogenous circadian timekeeping. To test this model, we have specifically abrogated these sites of daily neuronal remodeling through the reprogramming of neural development and assessed the effects on circadian timekeeping and clock outputs. Remarkably, the loss of these sites has no measurable effects on endogenous circadian timekeeping or on any of the major output functions of the small ventral lateral neurons. Rather, their loss reduces sites of glutamatergic sensory neurotransmission that normally encodes naturalistic time-cues from the environment. These results support an alternative model: structural plasticity in critical clock neurons is the basis for proper integration of light and temperature and gates sensory inputs into circadian clock neuron networks.

## Introduction

The proper daily timing of sleep and activity is the product of two processes, endogenous circadian timekeeping and the daily resetting of circadian rhythms to local time (i.e., entrainment) (Roenneberg et al., 2003). The importance of these two processes for health are made clear by a growing body of evidence that post-industrial light and social environments result in weak and unstable circadian entrainment, leading to a loss of sleep, increased cancer risk, and metabolic derangement (Roenneberg and Merrow, 2016). The master circadian clock, which drives daily rhythms in sleep and activity, resides in small islands of brain tissue (Herzog, 2007) wherein connections among diverse neuron types ensure a robustness in circadian timekeeping that is lacking in peripheral tissues (Hastings et al., 2018). Such circadian timekeeping networks require inputs from sensory pathways to entrain to daily environmental rhythms (Golombek and Rosenstein, 2010). Understanding the network properties of circadian timekeeping and entrainment in the brain is a central challenge in chronobiology.

Critical neurons within master timekeeping networks in both insect and mammalian brains undergo striking daily changes in cellular morphology (reviewed by (Bosler et al., 2015) (Krzeptowski et al., 2018)). In *Drosophila* the small ventrolateral neurons (s-LN_v_s) undergo daily clock-controlled structural remodeling, displaying significantly more extensive and highly branched dorsomedial projections in the early day relative to the early night (Fernández et al., 2008), a rhythm driven by daily changes in the outgrowth and de-fasciculation of terminal arborizations (Petsakou et al., 2015) (Sivachenko et al., 2013) (Gorostiza et al., 2014). The s-LN_v_s are critical for circadian timekeeping and properly timed behavioral outputs in fly and produce Pigment Dispersing Factor (PDF), which is likewise required for robust circadian timekeeping (Helfrich-Förster, 1998) (Renn et al., 1999). The termini of the s-LN_v_ dorsal projections contain synaptic and dense core vesicles (Yasuyama and Meinertzhagen, 2010) and their daily structural changes occur among the neurites of s-LN_v_ output targets (Gorostiza et al., 2014; Yasuyama and Meinertzhagen, 2010). For these reasons the dorsal termini of the s-LN_v_s have long been considered to be the major sites of s-LN_v_ axonal output (Helfrich-Förster, 1998) (Yasuyama and Meinertzhagen, 2010) and their daily structural plasticity is generally assumed to be a mechanism of circadian clock output in the brain (Bosler et al., 2015).

Within the hypothalamic suprachiasmatic nuclei (SCN), the master clock of the mammalian brain, neurons expressing the neuropeptide vasoactive intestinal poly-peptide (VIP) support circadian rhythms in a manner remarkably similar to PDF expressing clock neurons of *Drosophila*. The loss of VIP or its receptor results in a syndrome of circadian phenotypes that are highly reminiscent of those accompanying the loss of PDF or its receptor in the fly(Aton et al., 2005; Colwell et al., 2003; Hyun et al., 2005; Lear et al., 2005; Mertens et al., 2005; Renn et al., 1999). The VIP expressing neurons of the SCN undergo marked daily changes in morphology, displaying increased glial coverage of somata and dendrites during the day (Becquet et al., 2008). The morphological changes exhibited by the VIP expressing neurons of the SCN are accompanied by daytime increases in synaptic inputs, including glutamatergic inputs from the retinohypothalamic tract, onto VIP neurons (Girardet et al., 2010). Furthermore, retino-recipient Calbindin-D28K expressing neurons in the hamster SCN display elaborate arborizations in the early subjective night compared to other times (LeSauter et al., 2009). Taken together, the work on morphological plasticity in the rat and hamster SCN suggests that it may serve to mediate the integration of sensory (i.e., light) input (Girardet et al., 2010), however the large numbers and heterogeneity of neurons composing the SCN make a mechanistic examination of the functional import of such plasticity difficult to address experimentally.

Circadian neuronal remodeling of the *Drosophila* s-LN_v_s requires a functional circadian clock (Fernández et al., 2008) and neuronal firing promotes daily structural changes in dorsal termini of the s-LN_v_s (Sivachenko et al., 2013). Nevertheless, structural plasticity persists when these neurons are electrically silenced, revealing an endogenous cellular program for circadian structural plasticity (Depetris-Chauvin et al., 2011). Daily s-LN_v_ remodeling is driven by daily rhythms in clock-controlled gene expression (Depetris-Chauvin et al., 2014; Gunawardhana and Hardin, 2017; Petsakou et al., 2015; Sivachenko et al., 2013) and is therefore considered an output of the molecular clock within these neurons. The daily extension and retraction displayed by the s-LN_v_s is promoted by clock-driven oscillations in Fasciclin 2 (Fas2) mediated fasciculation/de-fasciculation rhythms (Sivachenko et al., 2013), metalloproteinase expression rhythms (Depetris-Chauvin et al., 2014), and rhythmic modulation of Rho1 GTPase signaling that drives daily terminal outgrowth and retraction (Petsakou et al., 2015).

An increasing number of studies have reported manipulations of the s-LN_v_ dorsal termini that are accompanied by significant effects on circadian timekeeping and output (Cusumano et al., 2018; Depetris-Chauvin et al., 2014; Gunawardhana and Hardin, 2017; Petsakou et al., 2015; Sivachenko et al., 2013). For example, the overexpression of the clock-controlled transcription factor Mef2 results in both constitutively open/complex termini and in a significant reduction in the percentage of flies able to maintain endogenous circadian rhythms in activity (Sivachenko et al., 2013). Likewise, the overexpression of the Rho1 GTPase in LN_v_ neurons results in both constitutively simple/closed termini and a significant weakening of locomotor rhythms (Petsakou et al., 2015). However, manipulations that cause significant morphological changes in the dorsal termini but nevertheless fail to alter free-running circadian rhythms or clock outputs have also been reported (Cusumano et al., 2018; Depetris-Chauvin et al., 2014; Petsakou et al., 2015; Sivachenko et al., 2013). Thus, the functional significance of daily s-LN_v_ structural plasticity has not been unequivocally established.

Here we take advantage of the genetic malleability and relative simplicity of the *Drosophila* clock neuron network to examine the functional significance of sites of circadian neuronal remodeling in the s-LN_v_s. By manipulating a well characterized mechanism of neuronal path finding, we have specifically prevented the development of the s-LN_v_ dorsal termini and comprehensively assessed the effects of their loss on endogenous circadian timekeeping and phasing of clock output. We find that the PDF-mediated timekeeping and output functions of the s-LN_v_s remain unchanged in the absence of these plastic terminal arborizations. Rather, we find that these termini mediate sensory inputs and the proper integration of time-cues from the environment. These results provide clear evidence that the sites of daily structural remodeling mediate sensory input, integration, and entrainment within the circadian clock neuron network and suggest that daily structural plasticity likely shapes the responses of circadian clock neurons to temporal cues from the environment.

### The expression of Unc-5 specifically prevents the formation of the s-LN_v_ dorsal projection termini

Previous work investigating the relationship between s-LN_v_ structural plasticity and circadian timekeeping employed genetic manipulations that clamped the dorsal termini in constitutively open or closed configurations, typically through the up- or down-regulation of transcription factors or cell signaling pathways (e.g. (Sivachenko et al., 2013) (Petsakou et al., 2015)). Most such manipulations have resulted in significant deficits in circadian sleep/activity rhythms, consistent with the longstanding hypothesis that s-LN_v_ plasticity mediates circadian timekeeping and output. However, several manipulations that produce defects in s-LN_v_ arbor morphology and/or plasticity have failed to produce circadian output phenotypes (e.g., (Sivachenko et al., 2013) (Depetris-Chauvin et al., 2014)), suggesting that dorsal termini manipulations that have produced circadian phenotypes may have acted via effects that were independent of the terminal arbor phenotypes they produced. Thus, though there is a significant body of evidence linking the sites of s-LN_v_ structural plasticity to circadian timekeeping and output, the functional significance of such plasticity remains an open question. For this reason, we sought to disrupt the formation of these termini developmentally to test the prediction that the absence of the sites of s-LN_v_ plasticity would produce timekeeping phenotypes reminiscent of the loss of s-LN_v_s or their major circadian peptide output PDF.

The formation of s-LN_v_ dorsal termini requires a turn toward the midline of the dorsal protocerebrum and the de-fasciculation of s-LN_v_ dorsal projections into fine radiating processes (Figures 1A and 1B; (Helfrich-Förster, 1995; Helfrich-Förster, 1997)). We found that the over-expression of the repulsive netrin receptor Unc-5 in all PDF expressing neurons completely abrogated the terminal ramification of the s-LN_v_ dorsal projections (Figure 1 and S1A), most likely by preventing the normal developmental outgrowth of these termini toward the midline where netrin is secreted during the development of the embryonic nervous system (Keleman and Dickson, 2001). Unc-5 overexpressing s-LN_v_s displayed a severely simplified dorsal projection that lacked the normal radiation of the dorsal medial termini (Figure 1A-D), a phenotype reflected by significant reductions in both the length of the dorsal projections and the brain volume they innervate (Figures 1E-J). Unc-5 overexpression was also accompanied by modest de-fasciculation of the ascending dorsal projection of the s-LN_v_s (Figure 1A, bottom right and S1C). Unc-5 overexpression had no obvious additional effects on the anatomical features of the small LN_v_s nor did it modify the anatomy of the large LN_v_s (Figures 1A, and S1B). We conclude that the overexpression of Unc-5 specifically prevents the formation of the s-LN_v_ dorsal termini, the sites of daily remodeling in these critical clock neurons.

**Figure 1.**
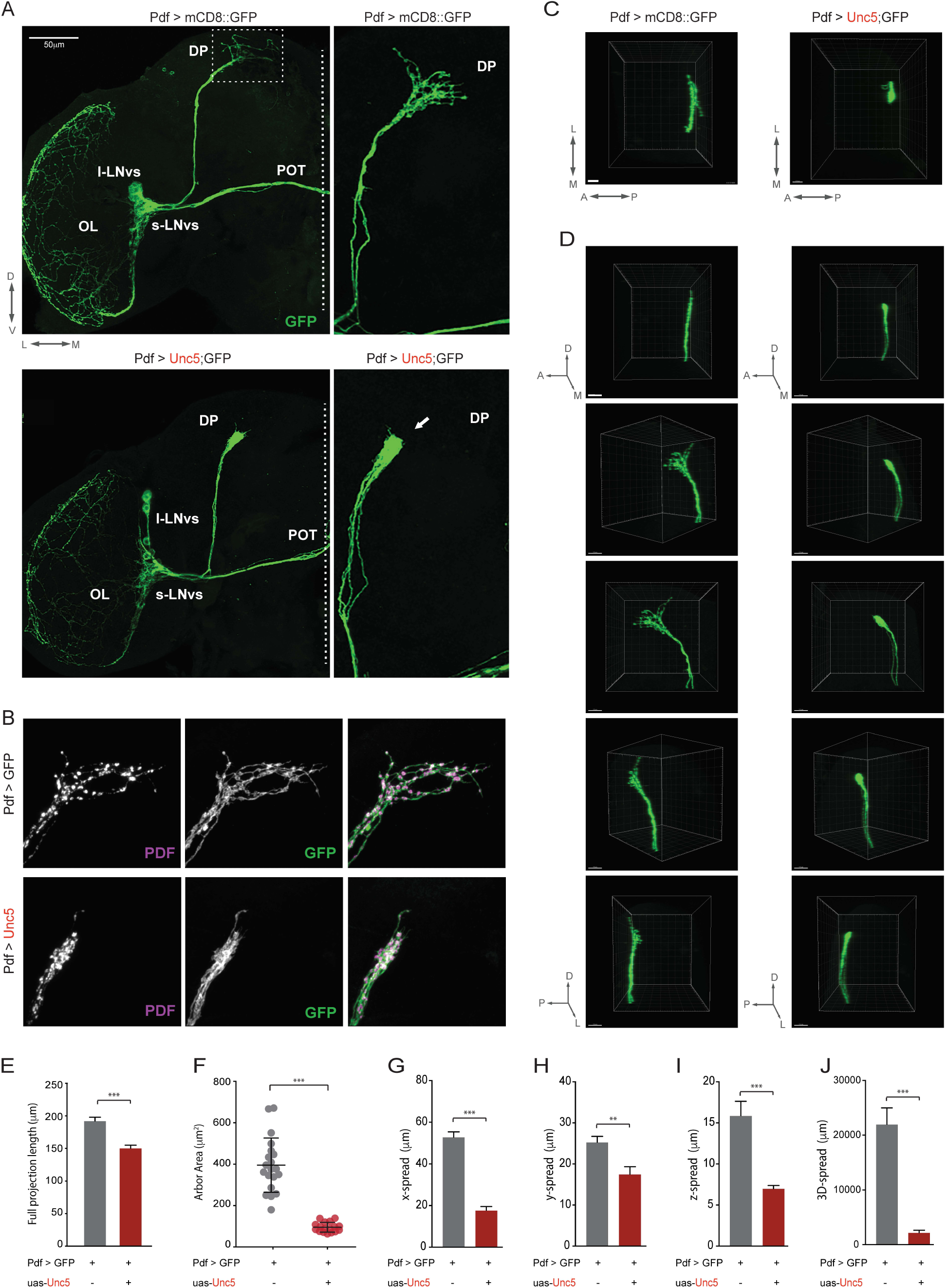
Overexpression of the axon guidance receptor Unc5 eliminates the dorsal arbor from the s-_v_s. (**A**) Representative confocal images of an anti-GFP immunostaining showing the left hemispheres of a ;*Pdf-Gal4/UAS-mCD8::GFP;* (top) and ;*Pdf-Gal4/UAS-mCD8::GFP;UAS-Unc5/+* (bottom) adult brain (left) and a magnified image of the s-LN_v_s projections extending into the dorsal protocerebrum (right). The cell bodies of the s-LN_v_s are located in the anterior ventral brain and their dorsal projections extend posteriorly and then turn toward the midline where they form a complex net of termini. Scale bar 50μm. Dorsal, ventral, medial, and lateral directions indicated at the bottom left of the top left panel. The dashed lines indicate the midline in the two left panels. l-LNvs indicate the large ventral lateral neurons, s-LNvs indicate the small ventral lateral neurons, DP indicates the dorsal projection, OL the medulla of the optic lobe, and OT, the posterior optic tract. (**B**) Confocal reconstructions of the dorsal termini of the s-LN_v_s in the dorsal protocerebrum of *Pdf-Gal4/UAS-mCD8::GFP;* (top) and ;*Pdf-Gal4/UAS-mCD8::GFP;UAS-Unc5/+* (bottom) flies. The left column shows anti-PDF immunostaining, the middle column anti-GFP immunostaining (green), and the right merged micrographs with PDF in magenta and GFP in green. (**C**) View of the dorsal termini of *Pdf-Gal4/UAS-mCD8::GFP;* (left) and ;*Pdf-Gal4/UAS-mCD8::GFP;UAS-Unc5/+* (right) dorsal termini through the dorsal surface of the brain. Scale bars = 15μM. (**D**) Comparison of 180 degree rotations of the dorsal projections of *Pdf-Gal4/UAS-mCD8::GFP;* (left) and ;*Pdf-Gal4/UAS-mCD8::GFP;UAS-Unc5/+* (right). Panels represent rotations of the projected Z-series starting from a lateral view and ending with a medial view of the projections. (**E-J**) Quantification of the effects of Unc5 expression on dorsal projection length (**E**), the brain area (i.e., X-Y spread) innervated by the dorsal termini in a collapsed Z-series from a posterior aspect (**F**) and the spread of these termini in the lateral medial (**G**), dorsal ventral (**H**) and anterior posterior (**I**) axes. The total three-dimensional spread of the dorsal termini is compared in (**J**). For E-J asterisks indicate significant differences. ** P < 0.01, *** P < 0.001. Error bars represent the standard error of the mean (SEM). See Table S1 for statistical information and sample sizes.

### PDF-mediated output functions of the s-LN_v_s do not require their dorsal termini

If the plastic dorsal termini of the s-LN_v_s are critical for circadian timekeeping and output signals, the loss of these termini should behaviorally phenocopy the ablation of these cells or the genetic loss of PDF, their major circadian output transmitter (Renn et al., 1999). The loss of the LN_v_s and PDF both result in a syndrome of timekeeping phenotypes that includes the loss of morning anticipation and an advance in the daily evening peak of activity under light/dark (LD) cycles and a significant weakening of the endogenous circadian rhythm under constant darkness and temperature (DD) accompanied by a decrease in free-running period (Renn et al., 1999)(Figures 2A, B, E and 3A). We first asked if the Unc-5 mediated prevention of dorsal termini development would be accompanied by phenotypes reminiscent of those caused by the loss of PDF under LD cycles. Under a 12h:12h LD cycle, the overexpression of Unc-5 in PDF expressing neurons had no measurable effects on the anticipation of LD transitions or on the entrained phase of evening peak activity, with *Pdf-Gal4/UAS-Unc5* flies displaying normal daily profiles of locomotion that that displayed the normal anticipation of light transitions (Figures 2C-E, S2, and S3). The s-LN_v_s exert control over much of the circadian clock neuron network through PDF mediated resetting signals (Stoleru et al., 2005) (Yao and Shafer, 2014) (Yao et al., 2016). When the molecular clocks within the s-LN_v_s are slowed down by the expression of the mutant clock kinase *Doubletime*^*LONG*^ (*Dbt*^*LONG*^), the daily evening peak of activity is delayed (Figure S4A), reflecting a resetting of the so called “evening cells” of the clock neuron network by PDF (Yao and Shafer, 2014) (Yao et al., 2016). Remarkably the s-LN_v_s were still able to set the evening peak of activity in the absence of their dorsal termini (Figure S4A).

**Figure 2.**
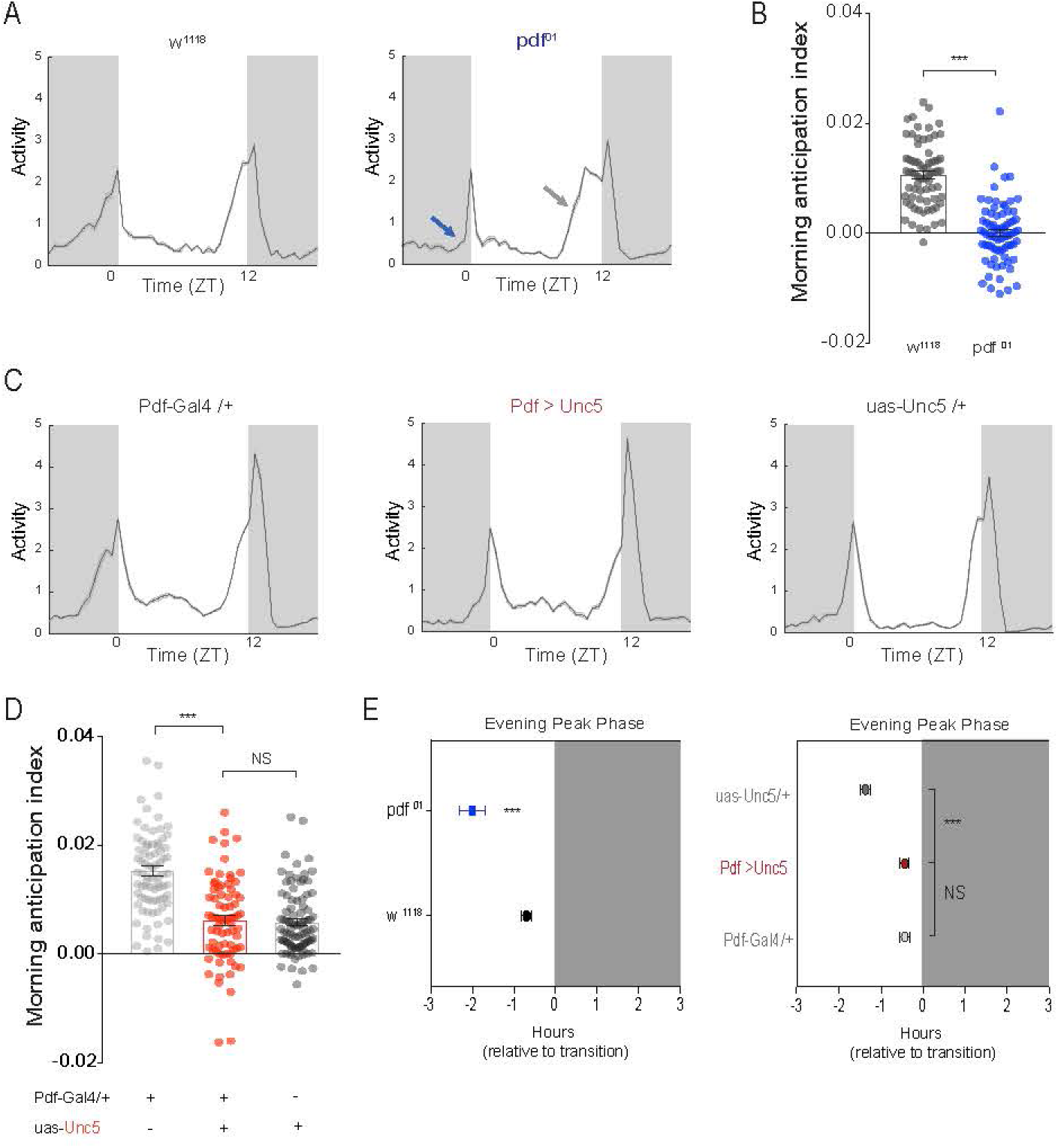
s-LN_v_ dorsal arbors are not required for PDF dependent behavioral outputs under light/dark conditions. (**A**) Population averaged activity profiles of wild-type (WT) and *Pdf*^*01*^ mutant flies during days two to six of entrainment to a 12h:12h LD cycle. Compared to the wild type activity profile (left panel) *Pdf*^*01*^ mutants lack morning anticipation (blue arrow) and exhibit an advanced evening peak of activity (gray arrow) (Renn et al., 1999). (**B**) The morning anticipation index is significantly different in WT and *Pdf*^*01*^ mutant flies. See also Figure S2A. (**C**) Population averaged activity profiles of *;Pdf-Gal4/+;UAS-Unc5/+* flies and their heterozygous parental controls reveal no *Pdf*^*01*^*-*like effects on morning or evening peaks of activity. (**D**) *;Pdf-Gal4/+;UAS-Unc5/+* flies do not differ significantly from their parental controls in morning anticipation (experimental flies are shown in red, dark gray indicates the *;;UAS-Unc5/+* parental control, light gray indicates the *;Pdf-Gal4/+;* parental control). (**E**) *;Pdf-Gal4/+;UAS-Unc5/+* flies also fail to display *Pdf*^*01*^ like evening peak phenotypes (WT and *Pdf*^*01*^ phases are shown on the left for comparison). Average evening peak phases are displayed +/- SEM. “0” marks the time of lights-off. Dark gray indicates night. See supplemental Table S1 for sample sizes and statistics.

The s-LN_v_s are the most dominant circadian pacemakers within the clock neuron network under conditions of constant darkness (DD) and temperature (Chatterjee et al., 2018). The loss of PDF peptide or genetic ablation of the LN_v_s dramatically weakens the endogenous circadian rhythm and produces a shortening of its free-running period under DD (Renn et al., 1999). Furthermore, when the speed of the molecular clock is changed within s-LN_v_s, PDF released from these neurons resets the molecular clocks within other clock neurons and thereby modulates the speed of systemic timekeeping (Stoleru et al., 2005) (Yao and Shafer, 2014) (Yao et al., 2016) (Chatterjee et al., 2018). If the plastic dorsal termini of the s-LN_v_s mediate these circadian output functions, we would expect to see clear timekeeping phenotypes under DD conditions. The loss of the dorsal termini in Unc-5 expressing LN_v_s was not accompanied by changes in the proportion of flies displaying endogenous circadian rhythms in locomotor activity, nor did it produce a shortening of its free-running period (Figures 3A-C, S4B, and Table 1). The expression of the mutant clock kinase *Dbt*^*LONG*^ only in the PDF expressing LN_v_s coherently sets the period of free-running sleep/activity rhythms to approximately 27 hours through a PDF-mediated resetting of the molecular clocks of target clock neurons (Yao and Shafer, 2014) (Yao et al., 2016). Remarkably, the co-expression of Unc-5 with *Dbt*^*LONG*^ in the LN_v_s did not prevent these neurons from lengthening the free-running period of locomotor rhythms or delaying the evening peak of activity on the first day of free-run ((Figure 3A, C, D, S4B and Table 1). Thus, the ability of the s-LN_v_s to control the clock neuron network was not affected by the absence of their normal sites of daily remodeling. We conclude that the normal sites of structural plasticity in the dorsal projections of the s-LN_v_s are not required for the established PDF-dependent output functions of these neurons.

**Table 1.**

| Genotype | Number of flies | % Rhythmic | Period ± SEM (h) | Rhythmic Power ± SEM |
| --- | --- | --- | --- | --- |
| *w<sup>1118</sup>* | 41 | 100 | 23.8 ± 0.04 | 137.5 ± 8.0 |
| *;Pdf-Gal4/+;* | 55 | 100 | 24.0 ± 0.02 | 172.4 ± 7.9 |
| *;;UAS-Unc5/+* | 62 | 98.4 | 23.6 ± 0.03 | 171.7 ± 6.2 |
| *;;UAS-Dbt<sup>L</sup>/+* | 58 | 100 | 23.9 ± 0.03 | 144.7 ± 5.4 |
| *;;UAS-Dbt<sup>L</sup> /UAS-Unc5* | 54 | 100 | 23.6 ± 0.03 | 152.9 ± 8.0 |
| *;Pdf-Gal4/+;UAS-Unc5/+* | 40 | 100 | 24.0 ± 0.04 | 134.1 ± 8.9 |
| *;Pdf-Gal4/+;UAS-Dbt<sup>L</sup>/+* | 36 | 96.6 | 27.2 ± 0.1 | 95.9 ± 9.6 |
| *;Pdf-Gal4/+;UAS-Dbt<sup>L</sup>/UAS-Unc5* | 58 | 100 | 26.2 ± 0.20 | 126.6 ± 7.7 |

**Figure 3.**
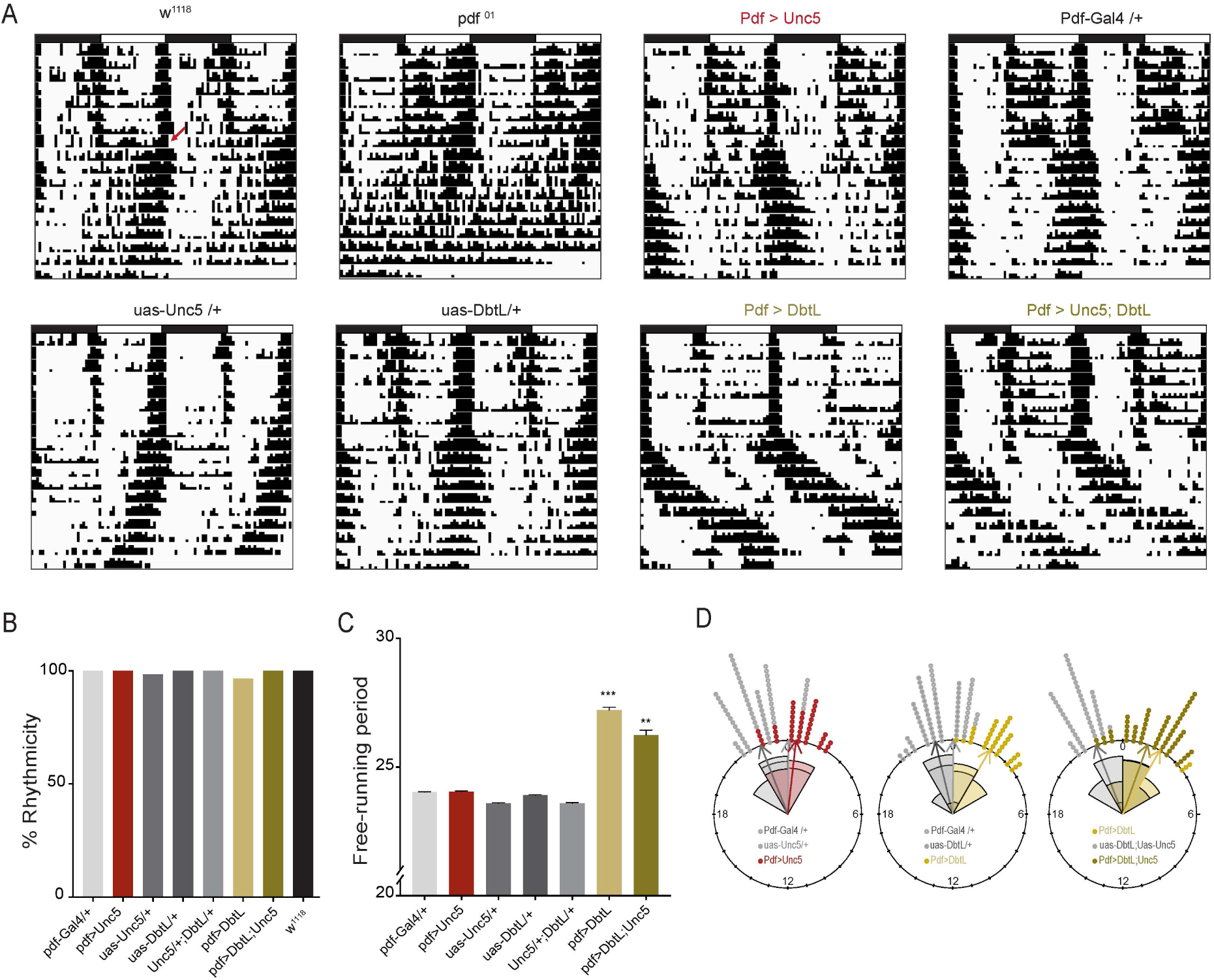
s-LN_v_s lacking dorsal termini maintain their control of systemic circadian timekeeping. (**A**) Representative double plotted actograms for flies under 8 days of LD entrainment followed by 10 days of free-running under constant darkness and temperature (DD) of the genotypes indicated. Both ;*Pdf-Gal4/+;UAS-Dbt*^*LONG*^*/+* and ;*Pdf-Gal4/+;UAS-Dbt*^*LONG*^*/UAS-Unc5/* flies exhibit significantly lengthened free-running periods. Red arrow indicates switch to DD conditions. (**B**) Summary of the percentage of flies displaying significant circadian periodicity under DD following entrainment to LD cycles. (**C**) Mean free-running period for seven days of DD activity rhythms. The endogenous periods of ;*Pdf-Gal4/+;UAS-Dbt*^*LONG*^*/+* and ;*Pdf-Gal4/+;UAS-Dbt*^*LONG*^*/UAS-Unc5/* flies are not significantly different and are significantly longer than all their parental control lines. ** P < 0.01, *** P < 0.001. Error bars indicate SEM. (**D**) Rose plots of evening activity peaks on the first day of free run under DD for the genotypes indicated. “0” marks the time 24-hours after the final lights-on event during the LD cycle. Control *;;uas-Unc5/+* flies displayed relatively early evening peak phases, but experimental *;Pdf-Gal4/+;uas-Unc5/+* flies did not differ from *;Pdf-Gal4/+* controls (left plot). The expression of *Dbt*^*LONG*^ in the Pdf-expressing neurons results in a significantly delayed evening peak (middle plot). The co-expression of *Unc5* with *Dbt*^*LONG*^ in the Pdf-expressing neurons did not prevent the delayed evening peak (right plot). Details of the circular statistical analysis are described in *Materials and Methods*. See also S1 for statistical information and sample sizes.

### The loss of s-LN_v_ dorsal termini causes deficits in the entrainment of locomotor rhythms to naturalistic ramping temperature cycles

The dorsal termini of the s-LN_v_s rest in close apposition to the neurites of the DN1_p_ class of clock neurons (Figure 4A-B), which are established targets of LN_v_ output (Shafer and Taghert, 2009; Zhang et al., 2010) and serve as major conduits of circadian output signals in the fly brain (Cavanaugh et al., 2014). Serial electron micrograph reconstructions of the s-LN_v_ termini revealed the presence not only of output (i.e., pre-synaptic) synapses, but also post-synaptic structures, indicating that the dorsal termini are axo-dendritic in nature (Yasuyama and Meinertzhagen, 2010). The DN1_p_s provide inhibitory feedback mediated by the release of glutamate onto the s-LN_v_s and to thereby promote sleep (Guo et al., 2016). Thus, the s-LN_v_s and DN1_p_s form bidirectional connections in the dorsal protocerebrum. The sites of PDF release from the dorsal termini of the s-LN_v_s appear to be extrasynaptic: PDF-containing dense core vesicles dock in regions of the dorsal projections that are not directly opposed by post-synaptic compartments (Yasuyama and Meinertzhagen, 2010), suggesting that PDF released from the dorsal termini may normally act at a distance. Unc-5 mediated abrogation of dorsal termini formation clearly reduced the volume of brain area through which s-LN_v_ and DN1_p_ neurites reside in apposition (Figure 4).

**Figure 4.**
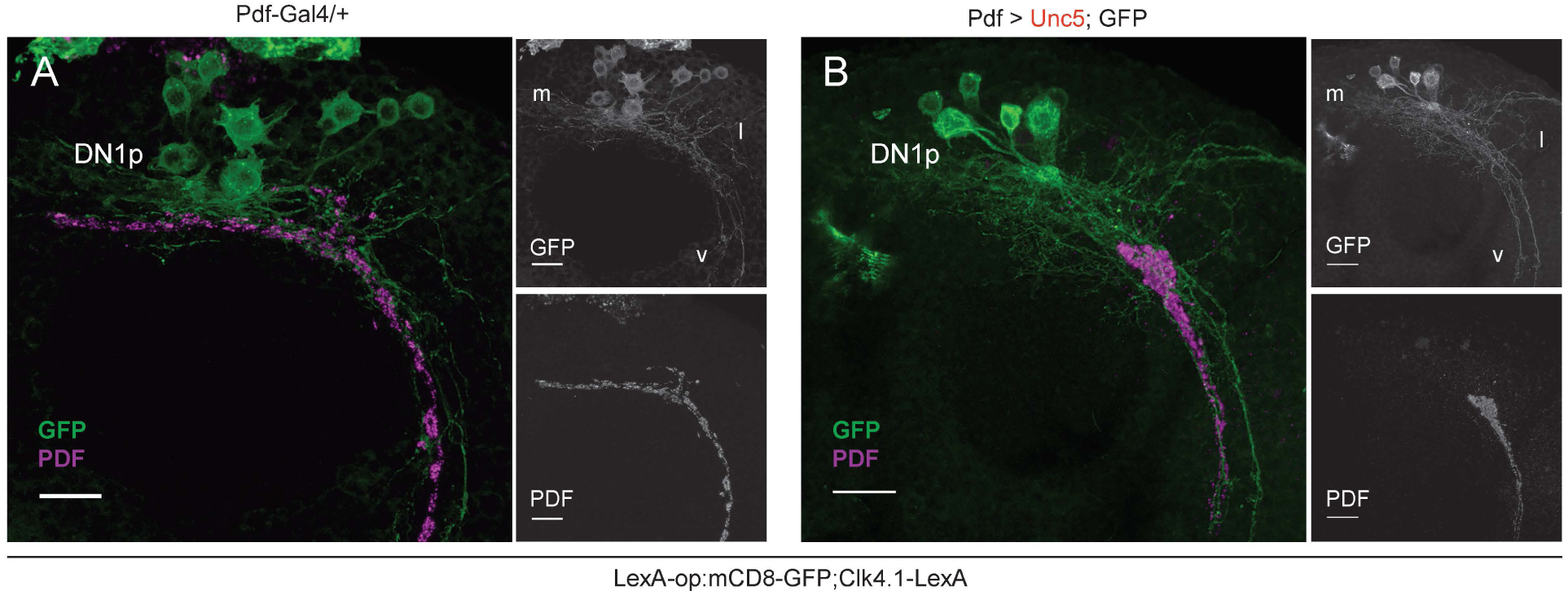
The expression of Unc5 causes significant changes in the anatomical relationship between the neurites of the DN1_p_s and the dorsal projections of the s-LN_v_s. (**A**) Confocal reconstruction of s-LN_v_ dorsal projections and the neurites of the DN1_p_ clock neurons in the dorsal protocerebrum of ;*Pdf-Gal4/LexAop-mCD8:GFP;Clk4.1LexA/+* flies. Brains were immuno-labelled for GFP (green) and PDF (magenta) and imaged through the posterior surface of the brain. Small panels display single gray scale reconstructions of GFP (top) and PDF expression (bottom). The medial (m), lateral (l) and ventral (v) extensions of the DN1_p_ neurons are indicated in the top right panel. (**B**) Confocal reconstruction of s-LN_v_ dorsal projections and the neurites of the DN1_p_ clock neurons in the dorsal protocerebrum of ;*Pdf-Gal4/LexAop-mCD8:GFP;Clk4.1LexA/UAS-Unc5* immunolabeled and imaged as described for **A**. Scale bars=30μm for all panels.

The DN1_p_s sensitively monitor environmental temperature (Yadlapalli et al., 2018) and their synaptic outputs are required for the normal entrainment of sleep/activity cycles to low amplitude step-function temperature cycles (Chen et al., 2015) and to gradually and constantly ramping temperature cycles (Yadlapalli et al., 2018). The organization of activity and sleep under such constantly changing temperature cycles is likely mediated by the inhibition of the s-LN_v_s by the DN1_p_s (Guo et al., 2016; Yadlapalli et al., 2018). Based on the close apposition of DN1_p_ neurites and s-LN_v_ dorsal termini and the axodendritic nature of the latter, we hypothesized that the plastic dorsal termini might be required for the normal entrainment to gradually ramping temperature cycles, predicting that the abrogation of the s-LN_v_ dorsal termini would lead to changes in the organization of sleep/activity rhythms under such entrainment conditions.

Under a 24-hour environmental temperature oscillation (20-28°C), which consisted of constant heating from 20 to 28°C for 12 hours followed by constant cooling from 28 to 20°C for 12 hours, *Pdf-Gal4/UAS-Unc5* flies differed significantly from their parental controls with regard to the daily pattern of activity. As previously described for wild type flies (Currie et al., 2009; Yadlapalli et al., 2018), the control *Pdf-Gal4/+* and *UAS-Unc5/+* flies displayed a rather small yet precipitous increase in activity at the onset of heating followed by gradual increases in locomotion throughout most of the heating phase and a precipitous drop in activity associated with the onset of cooling (Figure 5A and C). In contrast, *Pdf-Gal4/UAS-Unc5* flies displayed activity rhythms of significantly lower amplitude and did not begin their major daily increase in locomotion until the end of the heating phase, (Figures 5A and C). We quantified these features of entrainment in two ways: the ratio of activity levels seen near the end of the day (from ZT 10 to 12) to the magnitude of the small startle response at the onset of heating (from ZT00 to 02; Figure 5B) and the heating index (5D), which is based on the correlation between rising temperatures and locomotor activity (Figure 5D;(Yadlapalli et al., 2018)). Both metrics revealed significant differences between *Pdf-Gal4/UAS-Unc5* flies and their parental controls reflecting deficits in the ability to time daily activity increases with rising temperature. These results support the conclusion that the absence of s-LN_v_ dorsal termini was accompanied by an inability to properly entrain to naturalistic temperature cycles, likely due to the inability to integrate input from thermoreceptors via the DN1_p_s.

**Figure 5.**
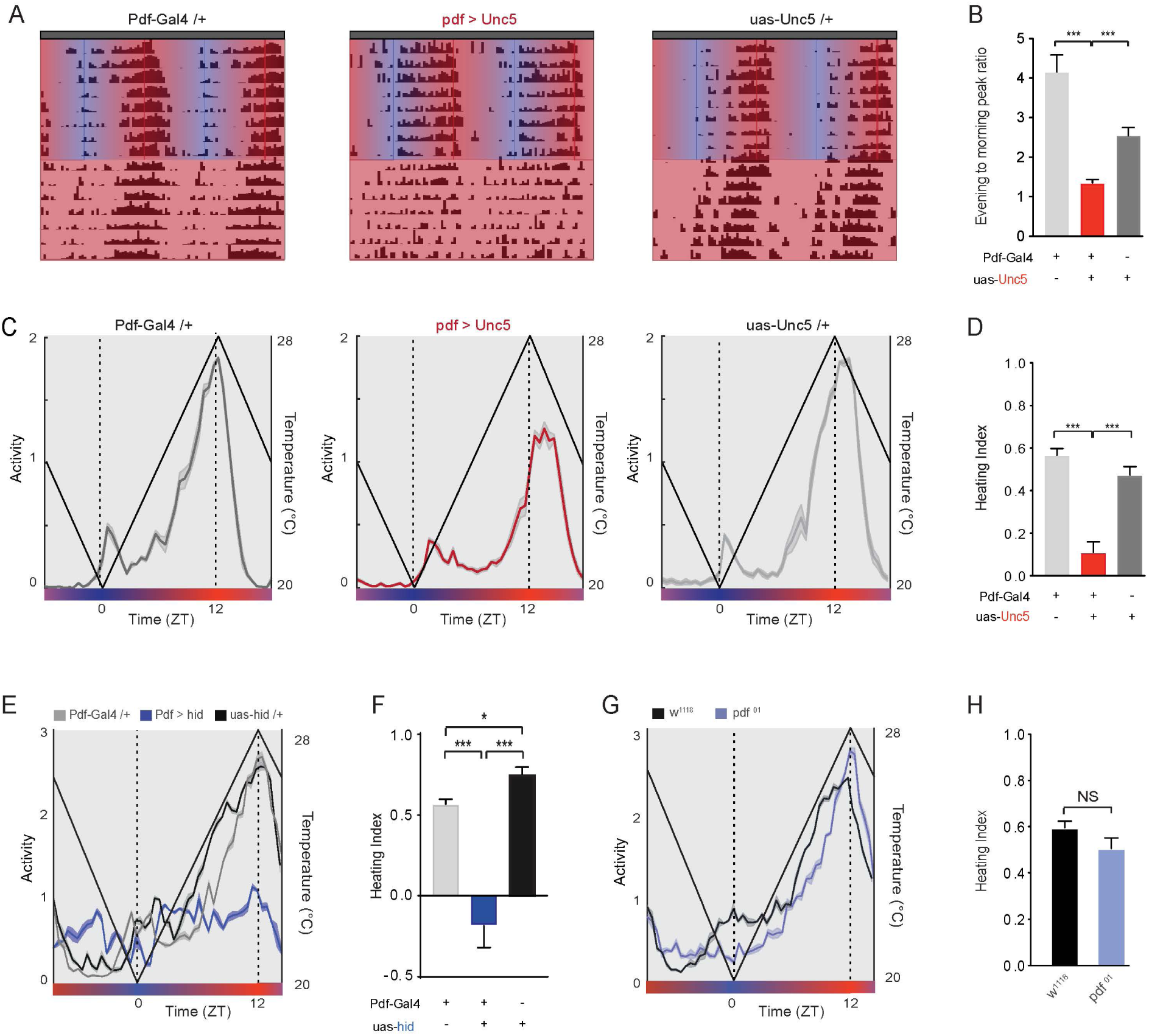
s-LN_v_ terminal arbors mediate entrainment to temperature ramps. (**A**) Representative actograms of single flies entrained for 8-days to constantly changing temperature ramps under DD followed by one week of free running at 25 °C under DD. During entrainment, temperature progressively increased from 20 °C to 28 °C between ZT 0-12 and gradually decreased from 28 °C to 20 °C between ZT 12-0. Blue to red gradients indicate heating phase, red to blue gradients indicate cooling phase. Genotypes are indicated above actograms. (**B**) Calculated ratios of evening peak activity between ZTs 10-12 to morning peak activity between ZTs 0-2 for flies of the following genotypes: *;Pdf-Gal4/+;, ;Pdf-Gal4/+;UAS-Unc5/+, and ;;UAS-Unc5/+.* (**C**) Averaged population activity plots for the last three days of entrainment to the temperature cycle (days six to eight). Straight black lines represent temperature changes. Dashed vertical lines indicate transition points between cooling and heating phases. Blue to red gradients indicate heating phase, red to blue gradients indicate cooling phase. ZT0 is the beginning of the heating phase (T= 20 °C), ZT12 is the end of the heating phase (T= 28 °C). (**D**) Heating indices, which reflect the correlation between environmental heating and increases in locomotor activity, for the genotypes indicated. (**E**) Averaged population activity plots and (**F**) heating indices for flies in which the proapoptotic gene *hid* was expressed in the PDF expressing LN_v_s compared to heterozygote parental controls. (**G**) Averaged population activity plots and (**H**) heating indices for *pdf* ^*01*^ mutants and their genetic background control, *w*^*1118*^. For all histograms, * P < 0.05, ** P < 0.01, *** P < 0.001, and NS indicates not significantly different. For all activity plots, lines represent mean ± SEM. See Table S1 for statistical information and sample sizes.

The genetic ablation of the PDF expressing LN_v_s results in a profound reduction in the amplitude of the activity rhythm under gradual temperature cycles, even more severe than those displayed by flies lacking dorsal termini: LN_v_-ablated flies displayed little evidence of locomotor rhythms under these conditions (Figure 5E-F). Remarkably, flies lacking PDF peptide displayed normal activity rhythms under such temperature cycles, coordinating their daily activity increases with rising temperature and displaying a precipitous decrease in activity associated with the onset of cooling (Figures 5G-H). We conclude that the normal entrainment of activity rhythms to constantly changing temperature cycles requires LN_v_ neurons but not the peptide transmitter PDF.

If the dorsal termini are important for the integration of temperature inputs, previous manipulations that abrogated the dorsal termini of the s-LN_v_s yet failed to produce circadian output phenotypes should produce clear behavioral phenotypes under gradual temperature cycles. We chose to examine flies over-expressing Fas2 in the LN_v_s. As previously described (Sivachenko et al., 2013), *Pdf-Gal4/UAS-Fas2;* flies display a profound and specific loss of the s-LN_v_ dorsal termini yet display completely normal activity rhythms under LD (Figure S5A-G) and strong normally-paced free-running activity rhythms under DD; (Figure S5 and S6A-B). However, under gradually ramping temperature cycles Fas2 induced loss of the s-LN_v_ dorsal termini produced phenotypes highly reminiscent of those displayed by flies whose s-LN_v_ termini development was prevented through Unc-5 expression (S6D-E). These results support the hypothesis that the dorsal termini of the s-LN_v_s are critical for the integration of temperature inputs for the entrainment of daily activity rhythms.

### The s-LN_v_ dorsal termini puncta display a sparse glutamate receptivity characterized by after-excitation

Excitation of the glutamatergic DN1_p_s produces inhibitory responses in the cell bodies and dorsal projections of the s-LN_v_s and bath applied glutamate causes hyperpolarization and Ca^2+^ decreases in s-LN_v_ cell bodies (Guo et al., 2016). Serial electron micrograph reconstruction of the s-LN_v_ dorsal projection revealed that their termini are sparsely dendritic, displaying approximately 10-fold fewer post-synaptic compared to presynaptic compartments (Yasuyama and Meinertzhagen, 2010), a ratio typical for axodendritic neurites in *Drosophila* (Takemura et al., 2008). If the dorsal termini of the s-LN_v_s mediate glutamate reception, we would therefore expect them to respond to bath applied glutamate directly and sparsely.

We characterized the effects of glutamate on individual puncta of the dorsal termini through the expression of the genetically encoded Ca^2+^ reporter GCaMP6f (Chen et al., 2013) (Figure 6A and D). Explanted brains from *Pdf-Gal4/+;UAS-GCaMP6f/+* flies were imaged in hemolymph-like saline containing 2µM tetrodotoxin (TTX) to inhibit the contribution of network influences and s-LN_v_ firing to the observed responses. The majority of puncta observed failed to display GCaMP6f responses to 30 second perfusion of 0.5 or 1mM glutamate (+TTX) (Figure 6A-C). However approximately 15% of optical sections revealed clear excitatory increases in GCaMP6f fluorescence among subsets of puncta immediately following wash-out or beginning shortly before the end of glutamate perfusion (Figure 6D-G). Thus, individual puncta of the s-LN_v_ dorsal projections appeared to be sparsely and directly receptive to glutamate, which caused a potent after excitation, suggestive of rebound excitation. Furthermore, these responses were often seen to begin just before the end of glutamate perfusion, suggesting desensitization. These features have been observed in the context of inhibition by ligand gated chloride channels (e.g., (Boehme et al., 2011; Gielen et al., 2015).

**Figure 6.**
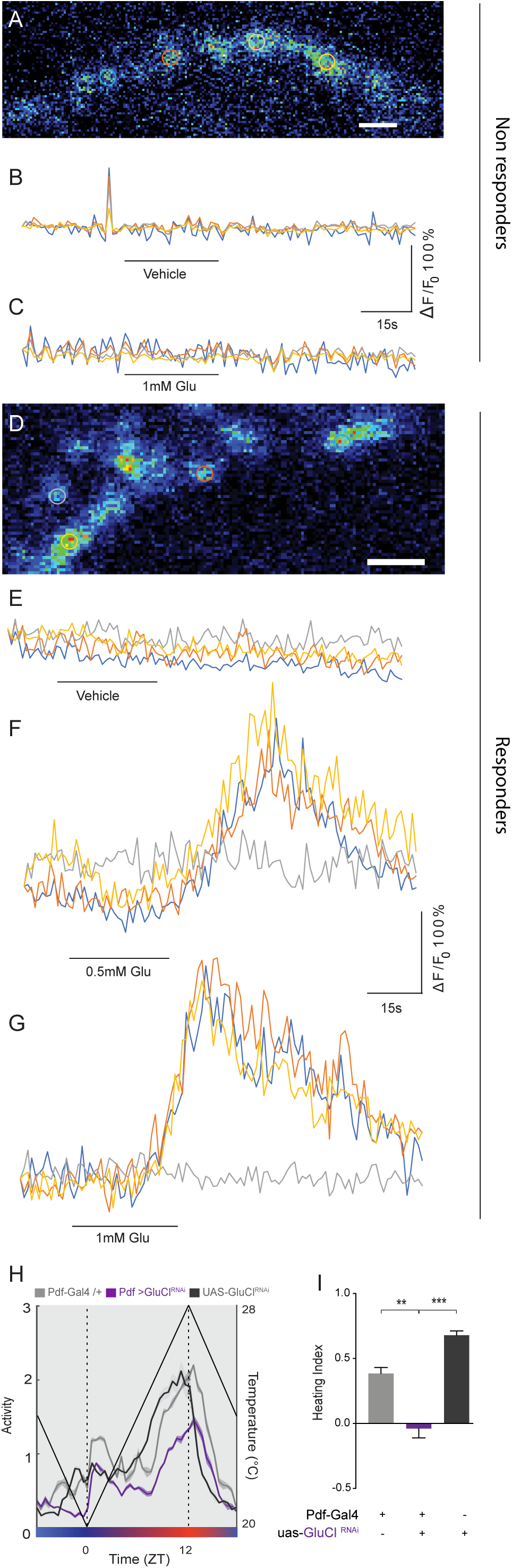
Puncta of the s-LN_v_ termini are sparsely receptive to glutamate and display rebound excitation following glutamate perfusion. **(A-C):** Representative glutamate responses for the majority of s-LN_v_ dorsal terminal puncta observed. (**A**) Expression of GCaMP6f in single puncta of the s-LN_v_ dorsal projections. Regions of interest (ROIs) are indicated for the plots below. Scale Bar = 5μm. (**B**) GCaMP6f fluorescence traces for the ROIs shown in **A** before, during, and after 30 s perfusion of vehicle (black bar). (**C**) GCaMP6f fluorescence traces for the ROIs shown in **A** before, during, and after 30 s perfusion of 1mM glutamate (black bar). (**D-G**). Representative glutamate response for the subset of glutamate receptive s-LN_v_ dorsal terminal puncta. (**D)** Expression of GCaMP6f in single puncta of s-LN_v_ dorsal projections. Regions of interest (ROIs) are indicated for the plots below. Scale Bar = 5μm. (**E**) GCaMP6f fluorescence traces for the ROIs shown in **D** before, during, and after 30 s perfusion of vehicle (black bar). (**F**) GCaMP6f fluorescence traces for the ROIs shown in **A** before, during, and after 30 s perfusion of 0.5mM glutamate (black bar), reveal a large excitatory response immediately after washout. GCaMP6f fluorescence traces for the ROIs shown in **A** before, during, and after 30 s perfusion of 1mM glutamate (black bar), reveal large excitatory responses that commence slightly before washout. Averaged population activity plots under ramping temperature cycles for experimental ;*Pdf-Gal4/UAS-GluClα-RNAi;UAS-Dicer2* flies (purple) and their parental heterozygote controls ;*Pdf-Gal4/+* (black) and ;*UAS-GluClα-RNAi/+;UAS-Dicer2/+* (gray). Plots represent the last three days of entrainment to a ramping temperature cycle (days 6-8), wherein temperature progressively increased from 20 °C to 28 °C between ZT 0-12 and gradually decreased from 28 °C to 20 °C between ZT 12-0. The straight black lines represent temperature changes. Dashed vertical lines indicate transition points between cooling and heating phases. Blue to red gradients indicate heating phase, red to blue gradients indicate cooling phase. ZT0 is the beginning of the heating phase (T= 20 °C), ZT12 is the end of the heating phase (T= 28 °C). (**I**) Heating indices for the genotypes shown in **H**, which reflect the correlation between environmental heating and increases in locomotor activity. ** P < 0.01; *** P < 0.001. See Table S1 for statistical information and sample sizes.

Regions of interest placed over the distal most regions of the of Unc-5 expressing s-LN_v_ dorsal projections (Figure S7A and D), which maintain the expression of the dendritic marker DscammTM2 ((Wang et al., 2004); Figure S7H-I), displayed a similar incidence of after excitation, though these responses typically appeared earlier during perfusion compared to normal dorsal termini. (Figure S7F-G). These results reveal that the puncta of the dorsal termini are normally sparsely receptive to glutamate and that the expression of Unc5 drastically changes the locations of s-LN_v_ dendritic processes in the dorsal protocerebrum.

### The knockdown of glutamate gated chloride channels in the s-LN_v_s results in temperature entrainment deficits

The glutamatergic DN1_p_s are thermo-receptive and required for the proper entrainment to environmental temperature cycles (Chen et al., 2015; Yadlapalli et al., 2018). Futhermore, these neurons form inhibitory connections onto the s-LN_v_s (Guo et al., 2016). We hypothesize that sites of daily remodeling in the s-LN_v_ dorsal termini are required for glutamatergic input from the DN1_p_ to the s-LN_v_ and that this input mediates the integration of temperature cycles into the circadian clock neuron network. We therefore predicted that the manipulation of glutamate receptors in the s-LN_v_s would result in significant changes in the organization of activity cycles under constantly changing temperature cycles that would be reminiscent of those associated with the absence of s-LN_v_ dorsal termini. The rebound excitation (Figure 6F-G) and apparent desensitization (Figure 6G) observed in our live imaging experiments were reminiscent of the behavior of cys-loop inhibitory receptors, which have been shown to produce rebound excitation in a manner dependent on voltage gated Ca^2+^ channels (Boehme et al., 2011) and whose mechanism of desensitization has been examined in detail (Gielen et al., 2015). For this reason, we examined the effects of the knockdown of the cys-loop glutamate gated chloride channels within the LN_v_s on the entrainment of activity rhythms to ramping temperature cycles.

The expression of RNAi constructs targeting the glutamate gated chloride channel GluClα in the LN_v_s resulted in phenotypes that were remarkably similar to those caused by the prevention of s-LN_v_ dorsal termini development (compare Figure 6H-I to Figures 5A-5D and S6C-E). The expression of GluClα-RNAi in LN_v_s resulted in lower amplitude activity rhythms and activity increases that failed to coincide with the daily rise in temperature (Figures 6H-I), further supporting the notion that the s-LN_v_s are critical for the reception of glutamate mediated temperature inputs. Thus, reducing the expression of GluClα in the s-LN_v_s phenocopied the loss of their dorsal terminal arbors, implicating these sites of structural plasticity in the reception of glutamatergic inputs relevant for the integration of temperature into the clock neuron network.

### The sites of s-LN_v_ plasticity are required for the integration of light and temperature by the circadian clock neuron network

Though the fruit fly readily entrains its sleep/activity rhythms to both light and temperature cycles, light has long been recognized as the most powerful environmental time cue for circadian entrainment (Zordan et al., 2001) (Roenneberg et al., 2013). However, temperature cycles can have strong effects on the timing of sleep and activity under LD cycles (Harper et al., 2016). Furthermore, the daily pattern of activity displayed by *Drosophila* in the field, wherein daily changes in both light and temperature occur, is significantly different from the bimodal pattern displayed by flies under the step function LD cycles and constant temperatures typically used in the lab, with flies in the field displaying a marked third peak of activity in the middle of the day (Vanin et al., 2012). The mechanisms through which the clock neuron network integrates light and temperature cues from the environment is not well understood. Our results reveal that the dorsal termini of the s-LN_v_ mediate the integration of temperature into the clock neuron network (Figures 5 and S6C-E). The s-LN_v_s are photoreceptive: they express the blue light sensor Cryptochrome (Yoshii et al., 2008) and receive synaptic inputs from external photoreceptors within the ventrally situated accessory medulla (Li et al., 2018; Schlichting et al., 2016). We therefore wondered if the dorsal termini of the s-LN_v_s might be critical for the integration of light and temperature cues from the environment.

In nature, diurnal temperature changes lag behind those of light due to heat exchange between air and the earth’s surface. To reflect this lag, we exposed flies to the standard 12:12 LD cycle and a constantly changing temperature cycle with the heating onset commencing one hour after lights on and the cooling onset commencing one hour after lights-off (Figure 7A and C, and S8). In wild-type (*w*^*1118*^), *Pdf-Gal4/+* and *UAS-Unc5/+* flies, the addition of the gradual temperature cycle resulted in a marked mid-day peak in activity, in addition to the expected morning and evening peaks under 12:12 LD, reminiscent of the midday peak of activity observed in the field but lower in amplitude (Figures 7A and C and S8; (Vanin et al., 2012)). Flies bearing Unc-5 mediated ablations of the s-LN_v_ dorsal termini failed to display the midday peak (Figures 7A-C), with *Pdf-Gal4/UAS-Unc5* flies displaying significantly lower midday activity than parental controls (Figure 7B). As for entrainment to gradual temperature cycles in DD (Figure 5G), the midday peak did not require PDF peptide (Figures 7D and S8). We conclude that the s-LN_v_ dorsal termini are required for the proper integration of light and temperature cues and normally serve as important sites for sensory integration within the *Drosophila* clock neuron network.

**Figure 7.**
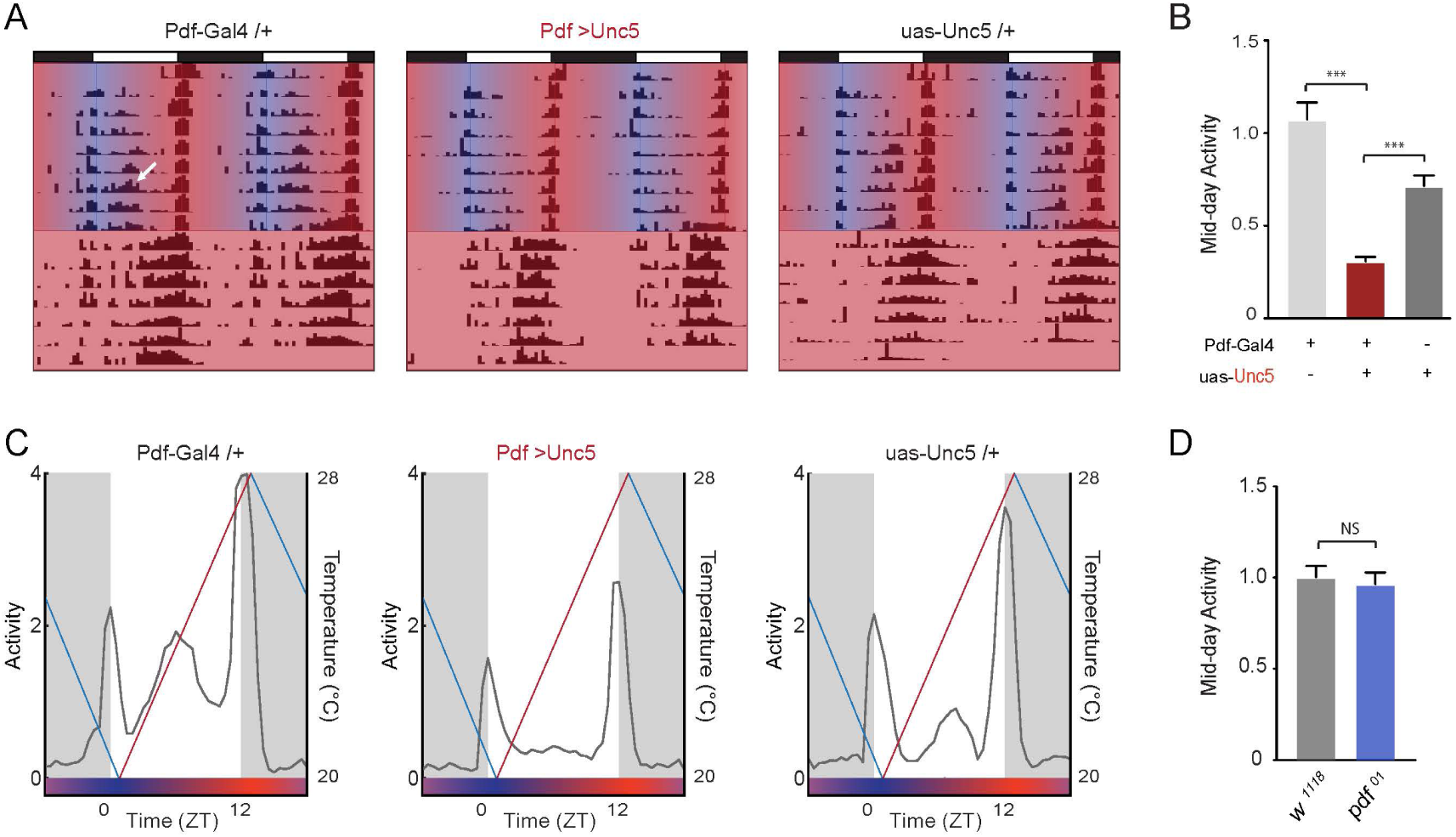
Prevention of s-LN_v_ terminal arbor development prevents the integration of temperature and light cycles. (**A**) Representative actograms of single flies *;Pdf-Gal4/+;, ;Pdf-Gal4/+;UAS-Unc5/+, and ;;UAS-Unc5/+* flies entrained for 8-days to constantly changing temperature ramps under a 12:12 LD cycle followed by one week of free running conditions under DD at 25 °C. During entrainment, temperature progressively increased from 20 °C to 28 °C between ZT 1-13 and gradually decreased from 28 °C to 20 °C between ZT 13-1, with heat and cooling commencing one hour after lights-on and lights-off, respectively. Blue to red gradients indicate heating phase, red to blue gradients indicate cooling phase. White arrow indicates a mid-day peak. (**B**) Comparison of mid-day activity levels for the genotypes shown in A. *** P < 0.001. (**C**) Averaged population activity plots of *;Pdf-Gal4/+;, ;Pdf-Gal4/+;uas-Unc5/+, and ;;UAS-Unc5/+* flies for five days (days two to six) of entrainment to offset ramping temperature cycle under LD. Straight red lines represent temperature increases, straight blue lines represent temperature decreases. Blue to red gradients indicate heating phase, red to blue gradients indicate cooling phase. ZT0 corresponds to lights-on, with heating commencing one hour after dawn. The presence of the temperature cycle has produced marked mid-day peaks in *;Pdf-Gal4/+;, and ;;UAS-Unc5/+* controls but fails to do so in experimental *;Pdf-Gal4/+;uas-Unc5/+* flies. (**D**) Comparison of mid-day activity in control (*w*^*1118*^) and *Pdf*^*01*^ mutant flies under a 12:12 LD cycle with an offset ramping temperature cycle. The loss of PDF peptide does not prevent the promotion of mid-day activity by ramping temperature cycles (see also Figure S7).

## Discussion

### The sites of daily remodeling in the s-LN_v_s are not required for circadian timekeeping or clock output function

A significant body of anatomical and genetic evidence supports the longstanding and widely accepted conclusion that the dorsal terminal arbors of these cells are critical sites of circadian output within the fly’s timekeeping network (e.g., (Helfrich-Förster, 1998) (Yasuyama and Meinertzhagen, 2010)). Surprisingly, we find that flies in which the development of these terminal ramifications has been prevented failed to display changes in PDF-mediated timekeeping or output functions. Furthermore, the absence of these dorsal termini did not prevent the s-LN_v_s from exerting their normal control over systemic circadian timekeeping under LD or DD conditions. These unexpected results lead us to conclude that the daily remodeling observed for these neurons is unlikely to mediate circadian output functions. Rather, they suggest that PDF mediated circadian output from the s-LN_v_s either acts over relatively long distances within the dorsal protocerebrum or that the major sites of circadian output take place in another region of the brain. Indeed, serial electron micrograph reconstructions of the s-LN_v_ dorsal projections reveal that PDF is released extrasynaptically in regions unopposed by post-synaptic regions of neighboring neurons (Yasuyama and Meinertzhagen, 2010). Furthermore, recent work has suggested that the accessory medulla of the ventral brain is an important site of PDF mediated circadian output (Schlichting et al., 2016). The presence of output synapses in the terminal arbors of the s-LN_v_s reveals that neurotransmitters are released by these plastic neurites. However, our results suggest that these outputs mediate signals that are not required for normal endogenous circadian timekeeping.

There is precedent for the timekeeping and output functions of the circadian system operating in the absence of synaptic connections with the nervous system. For example, the loss of wheel running rhythms caused by the bilateral ablation of the suprachiasmatic nuclei (SCN) is rescued by the implantation of fetal in the third ventricle of the brain (Lehman et al., 1987), even when the implant is encased in a semi-permeable capsule that prevents the outgrowth of neurons from the implant (Silver et al., 1996). Furthermore, genetically compromised cultured SCN slices that are characterized by arrhythmic expression of Period-Luciferase are rapidly rendered rhythmic when a functional SCN slice is placed in culture, with a period matching the circadian rhythm of the functional slice (Maywood et al., 2011). Rescue of clock gene cycling was achieved even when the slices were separated by a layer of mesh that prevented direct contact of the two slices while allowing for peptidergic communication (Maywood et al., 2011). In the context of these striking findings it is perhaps not surprising that specific fine ramifications of a particular group of clock neurons are not required for robust and properly timed circadian rhythms.

### The loss of the sites of daily remodeling in the s-LN_v_s prevents the integration of glutamate mediated time-cues

The dorsal projections of the s-LN_v_s are axodendritic (Yasuyama and Meinertzhagen, 2010) and s-LN_v_s appear to form bi-directional connections with the DN1_p_ class of clock neurons, cells that link the s-LN_v_s to neuroendocrine output pathways (Cavanaugh et al., 2014) and that provide feedback to the s-LN_v_s via glutamate-mediated inhibitory connections (Guo et al., 2016). DN1_p_ mediated inhibition of the s-LN_v_s appears to be critical for the entrainment of the sleep/wake cycle to constantly ramping temperature cycles (Yadlapalli et al., 2018). Our results indicate the plastic dorsal termini of the s-LN_v_s are necessary for the normal entrainment of the circadian clock to such temperature cycles and suggest that the puncta of these termini are sparsely receptive to glutamate via the cys-loop ligand gated chloride channel GluClα. We suggest that preventing the development of dorsal termini prevented the normal formation of DN1_p_ glutamatergic synapses onto the s-LN_v_ dorsal termini, leading to deficits in the integration of temperature inputs into the circadian clock neuron network. Thus, the sites of structural plasticity in the s-LN_v_s mediate sensory input and integration in the key set of clock neurons and daily structural changes in these termini likely result in changes in the number of inhibitory synapses between the DN1_p_s and the dorsal termini of the s-LN_v_s.

### Neuronal plasticity likely characterizes sensory input pathways in both insects and mammals

Neurons within the suprachiasmatic nuclei of the hypothalamus also display marked daily structural changes (LeSauter et al., 2009) (Girardet et al., 2010). Remarkably, the density of glutamatergic synapses onto VIP expressing neurons, cells that mediate functions strikingly similar to those of PDF expressing LN_v_s in the fly, were found to vary across the diurnal cycle (Girardet et al., 2010). Though the circadian functions of such remodeling have not been determined experimentally for mammals, they are hypothesized to underlie the entrainment of the clock to light/dark cycles (Girardet et al., 2010). Our work strongly links the sites of daily remodeling in a critical set of clock neurons in the fly with glutamate mediated input and the integration of environmental time-cues.

A canonical property of circadian rhythms is that the effect of environmental perturbation on the free-running system depends on the time at which it is delivered. The same perturbation delivered at various times in the circadian cycle can produce advances, delays, or have no effect on the subsequent phase of the rhythm (e.g., (De Coursey, 1960)). Given our findings that sites of structural plasticity mediate sensory input into the *Drosophila* clock neurons network, we hypothesize that daily changes in micro-anatomical features of clock containing neurons underlie the gating of such input into the clock neuron networks of both mammals and insects.

## Materials and Methods

### Fly Strains

Flies were reared on cornmeal-sucrose-yeast media under a 12hr:12hr light:dark (LD) cycle at 25 °C for standard LD-DD experiments or under constant darkness at 25 °C for temperature ramp experiments. The following fly lines were used in this study: *;Pdf(BMRJ)-Gal4; ;;Pdf01* and *;UAS-hid/CyO* (Park et al., 2000; Renn et al., 1999), (provided by P. Taghert, Wash U Med. School), *yw*;*Pdf-LexA;* (Shang et al., 2008) (provided by M. Rosbash, Brandeis), ;;*Clk4.1M-LexA* (Cavanaugh et al., 2014) (provided by A. Seghal, UPENN); *w;UAS-CD8:GFP;* (Lee and Luo, 1999); *w;;UAS-Dicer-2* (Bloomington Stock Center #24651), ;*UAS-GluClα*^*RNAi*^; (Vienna *Drosophila* Resource Center ID 105754) (Dietzl et al., 2007) w;*UAS-Fas2; (Dr. V. Budnik, UMSS Med. School), w;;UAS-Unc5-HA (Barry Dickson, Janelia Farm) (Keleman and Dickson, 2001), UAS-Dscam-TM2-GFP; Pin/CyO*; (Wang et al., 2004), *;;UAS-Dbt*^*LONG*^ *myc(27MIC)/(TM3)* (Jeffrey Price, University of Missouri at Kansas City) (Muskus et al., 2007), *w;;20xUAS-GCamp6f* (Bloomington Stock Number 52869), and *w;LexAop-mCD8GFP;TM2/TM6B,Tb* (Bloomington Stock Number 66545).

### Immunocytochemistry

Immunostaining of whole-mount *Drosophila* adult brains was done as previously described (Fernández et al., 2008). Flies were entrained to 12:12 LD cycles at 25°C and dissected brains were fixed in 4% paraformaldehyde for 1 hour at room temperature, blocked with 3% normal goat serum for 1 hour at room temperature, incubated with primary antibodies at 4° C overnight, and rinsed in PBS + 0.3% Triton (PBS-TX). The following antibodies were used: mouse anti-PDF (1:500, Developmental Hybridoma Bank), guinea pig anti-PAP (1:500, provided by Paul Taghert, Wash. U. Med School), and rabbit anti-GFP (1:1000, Invitrogen A-6455). Brains were rinsed of primary five times for 15 minutes or more with high agitation tumbling in PBS-TX and then kept in secondary antibody cocktail at 4° C overnight or for 2 h at room temperature and then rinsed in PBS-TX again as for primary. Brains were rinsed three times in PBS, mounted on a poly-L-lysine coated cover slip, dehydrated/cleared in a graded glycerol series (30%, 50% and 70% glycerol in PBS, 5-min each), and then mounted between coverslip bridges in HardSet Vectashield Mounting Medium (Vector Laboratories, Burlingame, CA). All samples were imaged on an Olympus Fluoview 1000 laser-scanning confocal microscope using either a UplanSApo 20x/0.75 NA or a 60x/1.10 NA W, FUMFL N objective (Olympus, Center Valley, PA). The arbor area and projection length of the s-LN_v_s were quantified using the Fiji platform (Schindelin et al., 2012) in ImageJ (Schneider et al., 2012). The length of the dorsal projection was determined by a line drawn from the point at which the s-LN_v_s dorsal projection and the posterior optic tract of the l-LN_V_s bifurcate near the accessory medulla to the end of the shortest neurite in control flies or at the distal end of the ‘bundle’ at the dorsal termini of Unc5 or Fas2 expressing s-LN_v_s. Area was determined by tracing the perimeter of the entire arbor in a projected Z-series. Imaris (Oxford Instruments, Abingdon, UK) was used for three dimensional reconstructions of the dorsal termini that were the basis of the quantification of x-spread, y-spread, z-spread, and total 3-D arbor spread.

### Live Imaging

*w;Pdf(BMRJ)-Gal4/+;UAS-GCaMP6f/+* and *w;Pdf(BMRJ)-Gal4/+;UAS-GCaMP6f/UAS-Unc5-HA* flies were anesthetized on ice, immobilized with a minuten pin through the thorax onto a 35mM Sylgard dish, and dissected under ice cold hemolymph-like saline (HL3) consisting of (in mM): 70 NaCl, 5 KCl, 1.5 CaCl_2_, 20 MgCl_2_, 10 NaHCO_3_, 5 trehalose, 115 sucrose, 5 HEPES; pH 7.1 (Stewart et al., 1994) containing 2μM Tetrodotoxin citrate (TTX) (Tocris, Bristol, U.K.). After dissection of all cuticle and pigmented eye tissue, brains were allowed to adhere to the bottom of poly-lysine coated 35 mm cellular culture dish (Becton Dickenson Labware, Franklin Lakes, NJ) under a drop of HL3 contained within a petri dish perfusion insert placed on the bottom of the dish with double sided adhesive (Bioscience Tools, San Diego, CA). Perfusion flow was established over the brain with a gravity-fed PS-8H perfusion system (Bioscience Tools, San Diego, CA). Test compounds were delivered to mounted brains by switching perfusion flow from the main HL3+TTX line to another channel containing 0.5 mM or 1.0 mM L-glutamate (Sigma Aldrich, St. Louis) in HL3 containing 2μM TTX, pH 7.1. To control for the effects of switching channels, we perfused HL3 + TTX from a second line as a vehicle control.

Live imaging was performed on an Olympus FV1000 laser-scanning microscope (Olympus, Center Valley, PA) under a 60x/1.10 NA W, FUMFL N objective (Olympus, Center Valley, PA). Single optical sections were scanned with a 488 nm laser at 1 Hz for 105 frames and GCaMP6f emission was directed to a photomultiplier tube by means of a DM405/488 dichroic mirror. Termini were imaged for 30s under constant HL3-TTX flow and then switched manually to a line containing 0.5 mM or 1.0 mM glutamate in HL3+TXX or a second line of HL3-TTX (Vehicle) for 15 or 30s and then switched back to the initial HL3-TTX line for the remaining frames of the time-course. For each dissected brain, vehicle controls were performed first followed immediately by glutamate treatments, starting with the lowest concentration. Time-courses characterized by significant movement artefacts during or after line switches were omitted from our analysis.

Time-courses of GCaMP6f fluorescence were measured in Fiji (Schindelin et al., 2012) in ImageJ (Schneider et al., 2012). Six to ten regions of interest (ROIs) were selected over single GCaMP6f expressing puncta of the s-LN_v_ dorsal termini, or over the bulbous ends of the Unc5-expressing dorsal projections, in which case six to ten ROIs of sizes comparable to those used over single puncta in normal dorsal termini. Mean pixel intensities (values between 0 and 4095) were collected for each ROI at each time point. Raw intensity plots were visualized for each ROI and plots were normalized to the initial fluorescence were constructed using the timepoint 15s before line switching as F_0_. Normalized plots were used to pool and compare vehicle and glutamate responses.

### Locomotor activity rhythm recording and analysis

Locomotor activity rhythms of adult male flies were recorded using DAM2 *Drosophila* Activity Monitors (TriKinetics, Waltham, MA). Three- to five-day old flies were placed individually in Trikinetics capillary tubes containing 2% agar-4% sucrose food at one end sealed with paraffin wax, plugged with a small length of yarn, and loaded onto the DAM2 monitors for locomotor activity recording. For standard LD entrainment and transfer to constant darkness (DD) free-run experiments, flies were entrained to 12:12 LD cycles for at least five days, and then released into constant darkness (DD) for at least eight days, at a constant temperature of 25°C. Activity counts were collected in 1-minute bins that were subsequently summed into 30-minute bins for the time-series analysis of locomotor activity.

Averaged population activity profiles of specific genotypes in LD were generated in Matlab (MathWorks, Natick). First, activity levels were normalized for individual flies, by setting the average activity level for all 30-min bins across the last four days in LD equal to 1.0. Population averages of this normalized activity were then determined for each 30-min bin over the number of LD cycles indicated in the results and figure legends. Finally, the population averages for the LD cycles were averaged into a single representative 24-hour day, which are displayed as either histograms or line plots.

Morning anticipation of light transitions under 12:12 LD was quantified by fitting 30-min binned beam crossing data over the last six hours of the night, with a least-squares linear regression. The beam crossing data for this six-hour window was averaged for the last 3 days of LD for individual flies and then normalized relative total activity for each fly within this window. These data were plotted for single flies in Matlab using the ‘scatter’ plotting function along with the least-squares regression lines fit to the average six-hour activity time-courses (Supplemental Figures S2 and S3). These scatter plots and regressions were overlaid with a line representing the average of the all individual fly regression lines. The slopes of individual regression lines were used as a metric of morning anticipation for single flies. The same approach was applied to the six hours preceding lights-on for the quantification of evening anticipation.

The phases of morning and evening peaks of individual flies on day one of DD were determined as previously described (Yao et al., 2016). Briefly, individual time-courses of beam crossings/30min through the first day under DD were subjected to a zero-phase Butterworth filter to diminish oscillations with periods of less than 20 hours (Levine et al., 2002). The ‘Findpeaks’ function in the Signal Processing Toolbox of Matlab was used for each fly’s filtered activity plot to identify the morning and evening peaks of activity, and their corresponding phases expressed as Circadian Time (CT). The morning and evening peak phases of experimental genotypes were compared to those of their parental controls using a Kruskal-Wallis one-way ANOVA and Dunn’s multiple comparison test. A summary of all pairwise comparisons is listed in Supplementary Table S1. In the case of the *w;Pdf(BMRJ)-Gal4/+;UAS-Dbt*^*LONG*^*/UAS-Unc5 flies*, we compared the behavior of these flies to all of the relevant single P-element heterozygotes and all double heterozygote combinations.

For circular statistics and rose plots, we transformed the negative and positive phases into proper hours on the 00-24h time scale by taking all phases modulo 24, and then converting the proper hours into radians. The zero-hour ZT00 is set at 24h, or 2π radians. We then applied the Watson two-sample test to determine whether the phases for control and experimental lines are significantly different. Watson’s non-parametric two sample U 2 statistic provides a criterion to test whether two samples differ significantly from each other. We performed nine tests using both the *Watson-Wheeler Test for Homogeneity of Angles* and the *Watson’s Two-Sample Test of Homogeneity* from the circular R library, designed and implemented for analyzing circular data. For both tests, the null hypothesis is that the two samples of angles come from the same underlying population.

We analyzed free-running activity rhythms using ClockLab software from Actimetrics (Wilmette, IL) as previously described (Yao and Shafer, 2014). In brief, rhythmicity, rhythmic power, and free-running period of individual flies were analyzed using Clocklab’s χ-square periodogram function implemented in ClockLab, based on a confidence level of 0.01 (Sokolove and Bushell, 1978). For all genotypes tested, significant periodicities between 14 and 34 hours were considered. For individuals that displayed more than one periodicity with a peak over significance, only the highest amplitude period was used for the determination of average periods displayed in Table 1 and S1. For each peak in the χ-square periodogram, Clock Lab returns a “Power” value and a “Significance” value. As previously described (Pfeiffenberger et al., 2010) (Yao and Shafer, 2014), “Rhythmic Power” was calculated by subtracting the Significance value from the Power value of the predominant peak for every fly designated as rhythmic, and was considered “0” for flies that failed to display a peak periodicity above significance.

To examine entrainment to naturalistic, gradually ramping temperature cycles, flies were reared under constant darkness (DD) at 25°C and then entrained to temperature cycles that gradually and constantly increased from 20 °C to 28 °C from ZT 00 to ZT 12 and gradually and constantly decreased from 28 °C to 20 °C from ZT 12-00 under DD. Flies were entrained under such temperature cycles for eight days. Averaged individual population activity plots were constructed for the last three days of temperature entrainment. The tracking of daily activity with rising environmental temperature was quantified as a “Heating Index” as described previously (Yadlapalli et al., 2018). Under the temperature conditions used here, flies displayed a startle response at the onset of heating that was dwarfed by the daily peak of activity that coincided with the warmest daily temperatures. To further quantify the extent to which daily activity rose with increasing daily temperature, we computed the ratios of evening peak activity (beam crossings between ZT 10-12) to morning peak activity (beam crossings between ZT 0-2) for days 6-8 of temperature ramp entrainment.

## Acknowledgments

We thank Paul Taghert, Rae Silver, and Patrizia Casaccia for critical reading a draft of this manuscript. We also thank Paul Taghert, and Bing Ye for useful discussions about the work and Barry Dickson, Greg Bashaw, Paul Taghert, Michel Rosbash, Vivian Budnick, and the Bloomington *Drosophila* Stock Center for sharing fly lines. The authors would like to acknowledge the Live Imaging and Bioenergetics Facility of CUNY Advanced Science Research Center for instrument use, and scientific and technical assistance and its director Dr. Ye He for sharing her expertise and for her assistance in the use of Imaris software. We also thank the members of the Fernández and Shafer labs for comments on the text.

## Supplemental Figures

**Figure S1. Overexpression of the axon guidance receptor Unc5 specifically affects the s-LN**_**v**_ **dorsal termini.** (**A**) Confocal Z-series reconstructions of five examples of anti-GFP immunolabeling of brains from *Pdf-Gal4*/*UAS-mCD8::GFP;UAS-Unc5/+* flies revealing the extent to which the development of the dorsal termini of the of s-LN_v_ dorsal projection was prevented by Unc5 expression. Images represent an scanning area of 75 μm × 75 μm. All the brains examined (n=40) revealed a complete absence of the dorsal termini. (**B**) The *UAS-Unc5* element alone does not cause arbor phenotypes (left). The posterior optic tract (POT) of the large LN_v_s was not affected by the expression of Unc5 (right panel). Scale bar = 50 μm. (**C**) Unc5 expressing s-LNvs display a modest de-fasciculation of ascending dorsal projection, consistently displaying more visually distinct, un-fasciculated neurites than controls (see also Figure 1A, lower right panel). *** P < 0.001.

**Figure S2. Anticipation indices reflect activity before the lights on and off transitions.** (**A**) Least-squares linear regression of normalized 30-min binned activity levels of individual flies (gray points and lines) during the last six hours of the night. Slopes of the individual fly regressions were used to quantify morning anticipation. The averaged regression line is shown in red. As expected, both the *Pdf*^*01*^ mutant and the *Clk*^*jrk*^ mutant lack the gradual increase in activity seen in wild type flies in the hours before lights on. (**B**) Evening Anticipation Index: an equivalent six-hour analysis of activity during the six hours before the lights-off transition for the same flies shown in A. Least-squares linear regression of normalized 30-min binned activity levels of individual flies are indicated by the gray points and lines. The averaged regression line is shown in blue. While the *Clk*^*jrk*^ mutant lacks the gradual increase in evening activity seen in wild type flies, the *Pdf*^*01*^ mutant exhibits clear anticipation of lights-off.

**Figure S3. Neither morning nor evening anticipation are affected by Unc5 overexpression in Pdf+ cells.** (**A**) The mean morning peak phase of experimental *;Pdf-Gal4/+;UAS-Unc5/+* flies is not significantly different than that of *;;uas-Unc5/+ controls*. **(B**) *Pdf-Gal4/+;UAS-Unc5/+* flies display robust free-running rhythms of locomotor activity, indistinguishable from their parental controls. (**C**) The least-squares regression approach to the quantification of evening peak reveals robust anticipation in both wild-type (*w*^*1118*^) and *Pdf*^*01*^ mutant flies. (**D**) Evening anticipation indices were not significantly different between *;Pdf-Gal4/+;UAS-Unc5/+* experimental flies and *Pdf-Gal4/+* controls. *** P < 0.001 and NS indicates no significant difference between groups.

**Figure S4. Unc5 expression in the LN**_**v**_**s does not prevent a slow molecular clock from inducing a long free running period of activity rhythms.** (**A**) Population averaged activity profiles of ;;*UAS-Dbt*^*LONG*^*/+* controls (left), *Pdf-Gal4*/*UAS-Dbt*^*LONG*^ flies (center), and ;*Pdf-Gal4*/+;*UAS-Dbt*^*LONG*^*/UAS-Unc5* (right). The expression of Unc-5 did not prevent the resetting of the evening peak (arrows) by the *Pdf*-expressing LN_v_s. (**B**) Representative χ-square periodograms for flies under seven days of free-running conditions (DD). Genotypes are indicated above the periodograms. Both ;*Pdf-Gal4/+;UAS-Dbt*^*LONG*^/+ and ;*Pdf-Gal4/+;UAS-Dbt*^*LONG*^/*UAS-Unc5* flies exhibit significantly longer free-running periods compared to all parental controls. See Table S1 for statistical information and sample sizes.

**Figure S5. Fas2-mediated elimination the dorsal termini of the s-LN**_**v**_**s does not affect the timing of activity under LD cycles.** (**A-C**) Representative confocal images of an anti-GFP immunostaining showing the left hemisphere of a ;*Pdf-Gal4/+*;*UAS-mCD8::GFP,UAS-Fas2/+* adult brain (**A**) and a magnified image of the s-LN_v_ dorsal projection (**B**) top panel, anti-PDF staining middle panel, anti-GPF staining bottom panel, merged images with PDF shown in magenta and GFP shown in green. (**C**) Examples of the absence of s-LNv dorsal termini ramification in five brains from ;*Pdf-Gal4/+UAS-mCD8::GFP;UAS-Fas2/+* flies. Images represent an area of 75 μm x 75 μm. (**D**) Quantification of the length of the s-LN_v_s projection for control ;*Pdf-Gal4/UAS-mCD8::GFP;* and experimental *;Pdf-Gal4/UAS-mCD8::GFP;UAS-Fas2/+* brains. (**E**) Quantification of area of s-LN_v_s dorsal terminal innervation for the genotypes shown in D. (**F**) Population averaged activity plot for ;*Pdf-Gal4*;*/+;UAS-Fas2/+* flies during days 3-5 of a 12h:12h LD cycle at a constant 25 °C. Neither the morning nor the evening peak are affected by the expression of Fas2. (**G**) Morning anticipation indices for ;*Pdf-Gal4*;*/+;UAS-Fas2/+* (blue) and for ;*Pdf-Gal4*;*/+;* and ;*;UAS-Fas2/+* controls (gray). See Table S1 for sample sizes and statistical information. *** P < 0.001 and NS = Not Significant. Error bars represent SEM.

**Figure S6. Fas2-mediated elimination the dorsal termini of the s-LN**_**v**_**s does not affect endogenous circadian timekeeping but impairs entrainment to temperature ramps.** (**A**) Normalized activity during the first three days of free-running conditions under DD. Dark gray indicates subjective night and light gray indicates subjective day. (**B**) Percentage of rhythmic flies under DD, based on ten days of free-run. There were no significant differences between the three genotypes as determined by a Fisher’s exact test. (**C**) Averaged population activity plots for the last three days of entrainment to gradually ramping temperature cycles (days 6-8) for the genotypes indicated. The straight black line represents temperature change. Blue to red gradients indicate heating phase, red to blue gradients indicate cooling phase. ZT0 is the beginning of the heating phase (T= 20 °C), ZT12 is the end of the heating phase (T= 28 °C). (**D**) Representative actograms of single flies entrained for 8-days to constantly changing temperature ramps under DD followed by one week of free running at 25 °C under DD. During entrainment, temperature progressively increased from 20 °C to 28 °C between ZT 0-12 and gradually decreased from 28 °C to 20 °C between ZT 12-0. Blue to red gradients indicate heating phase, red to blue gradients indicate cooling phase. Genotypes are indicated above actograms. (**E**) Calculated ratios of evening peak activity between ZTs 10-12 to morning peak activity between ZTs 0-2 for flies of the following genotypes: *;Pdf-Gal4/+;, ;Pdf-Gal4/+;UAS-Fas2/+*, and *;;UAS-Fas2/+.* The heating index did not capture the clear differences in amplitude displayed by experimental flies in this case.

**Figure S7. Truncated Unc5 expressing s-LN**_**v**_ **termini are sparsely receptive to glutamate and display rebound excitation following glutamate perfusion. (A-C):** Representative glutamate responses for the majority of truncated s-LN_v_ dorsal termini observed from ;*Pdf-Gal4/+;UAS-GCaMP6f/UAS-Unc5* flies. (**A)** Expression of GCaMP6f in the dorsal terminus of Unc5-expressing s-LN_v_s. Regions of interest (ROIs) are indicated for the plots below. Scale bars = 5μm. (**B**) GCaMP6f fluorescence traces for the ROIs shown in **A** before, during, and after 30 s perfusion of vehicle (black bar). (**C**) GCaMP6f fluorescence traces for the ROIs shown in **A** before, during, and after 30s perfusion of 1mM glutamate (black bar). (**D-G**). Representative glutamate responses for receptive s-LN_v_ dorsal terminal puncta. (**D)** Expression of GCaMP6f in the dorsal terminus of Unc5-expressing s-LN_v_s. Regions of interest (ROIs) are indicated for the plots below. Scale bars = 5μm. (**E**) GCaMP6f fluorescence traces for the ROIs shown in **D** before, during, and after 30 s perfusion of vehicle (black bar). (**F**) GCaMP6f fluorescence traces for the ROIs shown in **A** before, during, and after 30 s perfusion of 0.5 mM glutamate (black bar), reveal a large excitatory response that commences slightly before washout. (**G**) GCaMP6f fluorescence traces for the ROIs shown in **A** before, during, and after 30 s perfusion of 1.0 mM glutamate (black bar), reveal a large excitatory response that commences well before washout. (**H**) Expression of the dendritic marker DscamTM2GFP in the s-LN_v_ dorsal projection ;*Pdf-Gal4/UAS-DscamTM2-GFP*; flies. Anti-PDF immunostaining is shown on the left, anti-GFP immunostaining is shown on the right. (**I**) Expression of the dendritic marker DscammTM2GFP in the s-LN_v_ dorsal projections from ;*Pdf-Gal4/UAS-DscamTM2-GFP;UAS-Unc5/+* flies. Anti-PDF immunostaining is shown on the left, anti-GFP immunostaining is shown on the right. The truncated dorsal termini of Unc5-expressing s-LN_v_s maintain the expression of this dendritic reporter.

**Figure S8. The production of a daily, temperature-induced mid-day peak under LD cycles does not require PDF.** (**A**) Averaged population activity plots of control *w*^*1118*^ flies and *Pdf*^*01*^ mutants for five days under offset temperature cycle under LD (days two to six). Straight red lines represent temperature increases, straight blue lines represent temperature decreases. Blue to red gradients indicate heating phase, red to blue gradients indicate cooling phase. ZT0 corresponds to lights-on, with heating commencing one hour after dawn. The presence of the temperature cycle results in marked mid-day peaks in both control (*w*^*1118*^) and *Pdf*^*01*^ mutant flies. (**B**) Representative actograms of control (*w*^*1118*^) and *Pdf*^*01*^ mutant flies entrained for 8-days to constantly changing temperature ramps under a 12:12 LD cycle followed by one week of free running conditions under DD at 25 °C. During entrainment, temperature progressively increased from 20 °C to 28 °C between ZT 1-13 and gradually decreased from 28 °C to 20 °C between ZT 13-1, with heat and cooling commencing one hour after lights-on and lights-off, respectively. Blue to red gradients indicate heating phase, red to blue gradients indicate cooling phase. White arrows indicate mid-day peaks.

