## Supplementary Figures and Table S1. for "Sites of Circadian Clock Neuron Plasticity Mediate Sensory Integration and Entrainment"

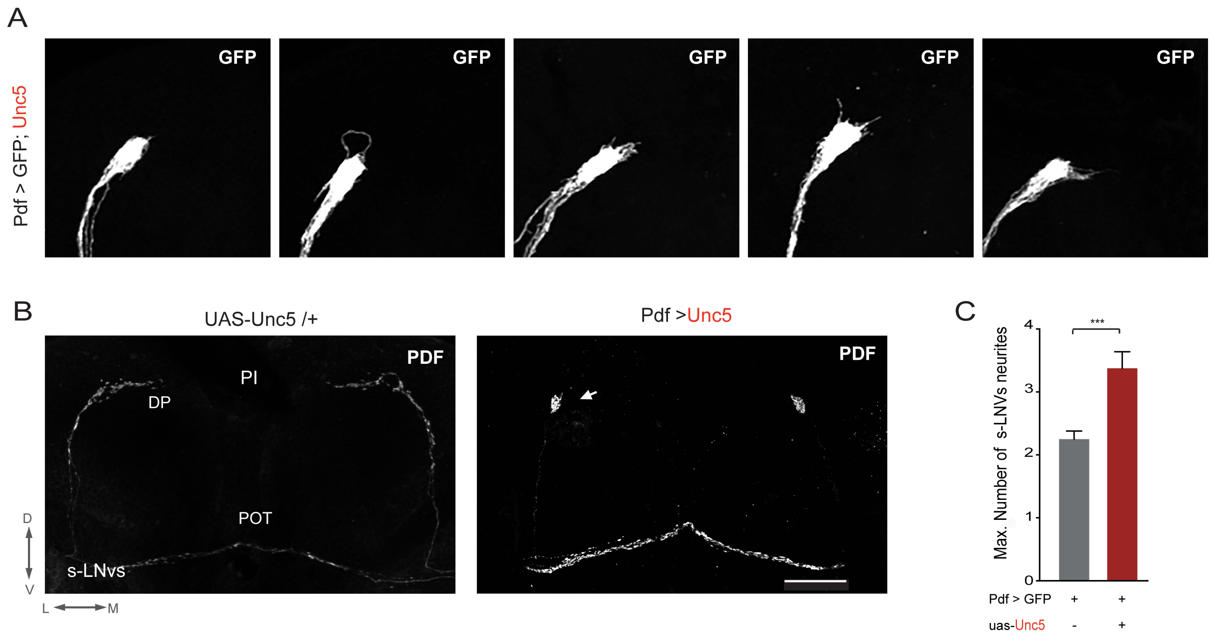

**Figure S1**. Overexpression of the axon guidance receptor Unc5 eliminates the dorsal arbor from the s-LNvs.

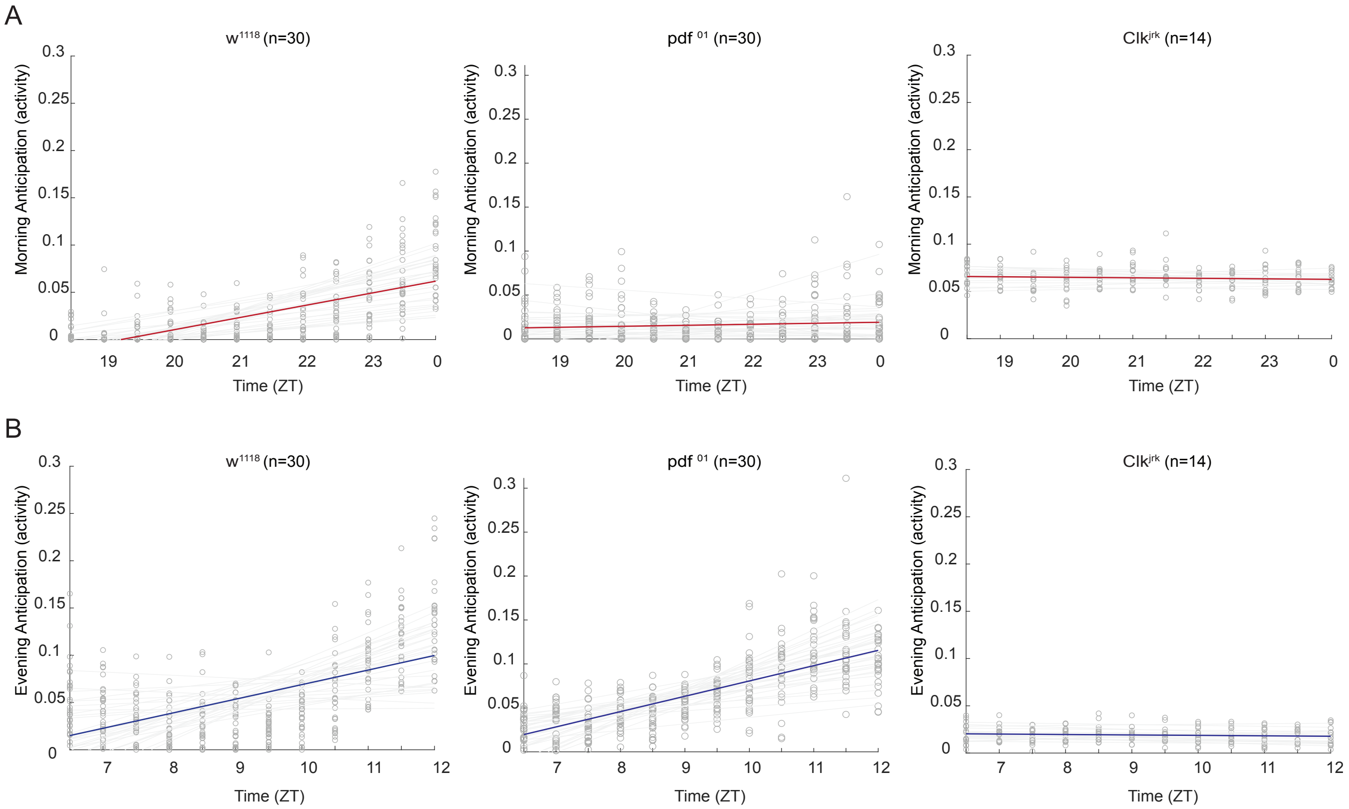

**Figure S2**. Anticipation indices reflect activity before the lights on and off transitions.

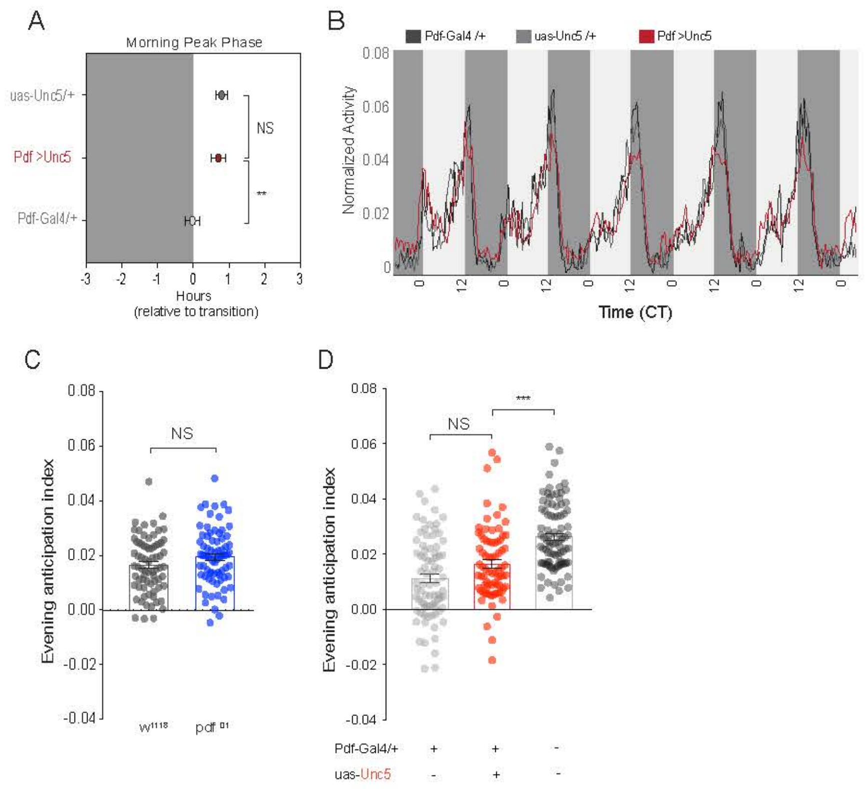

**Figure S3**. Neither morning nor evening anticipation are affected by Unc5 overexpression in Pdf+ cells.

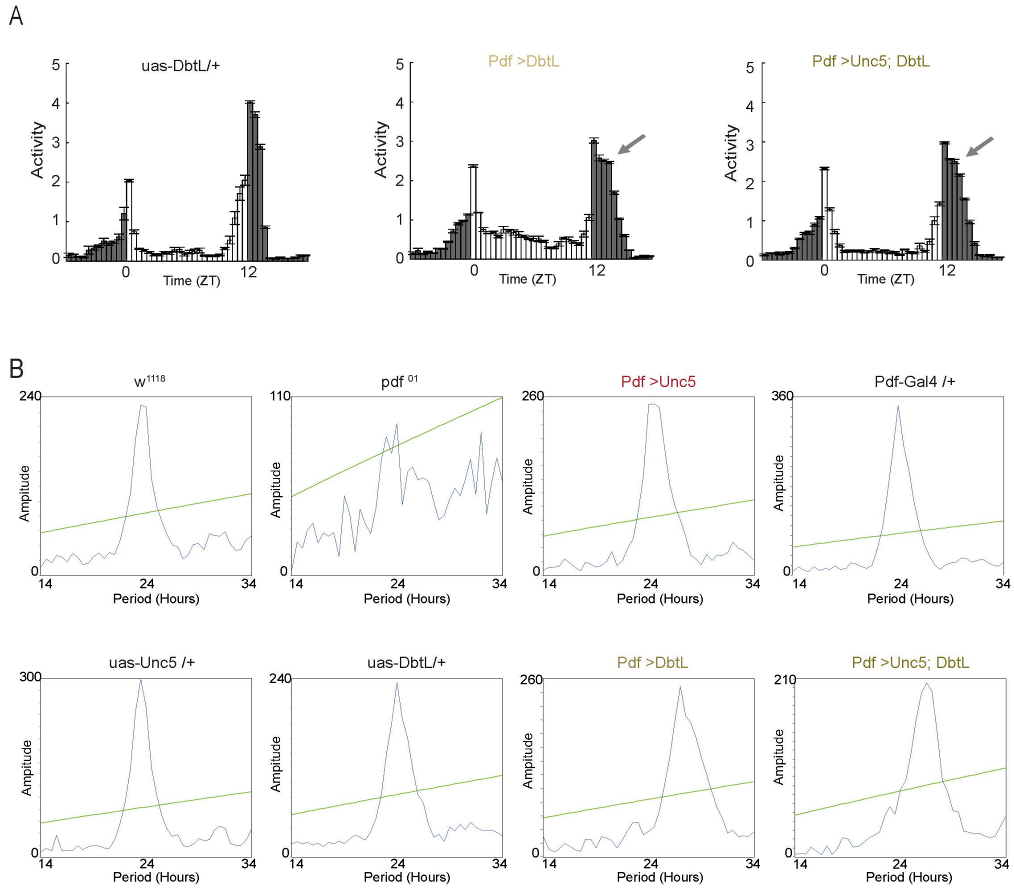

**Figure S4**. Unc5 expression in the LNvs does not prevent a slow running molecular clock from inducing a long free running period of activity rhythms.

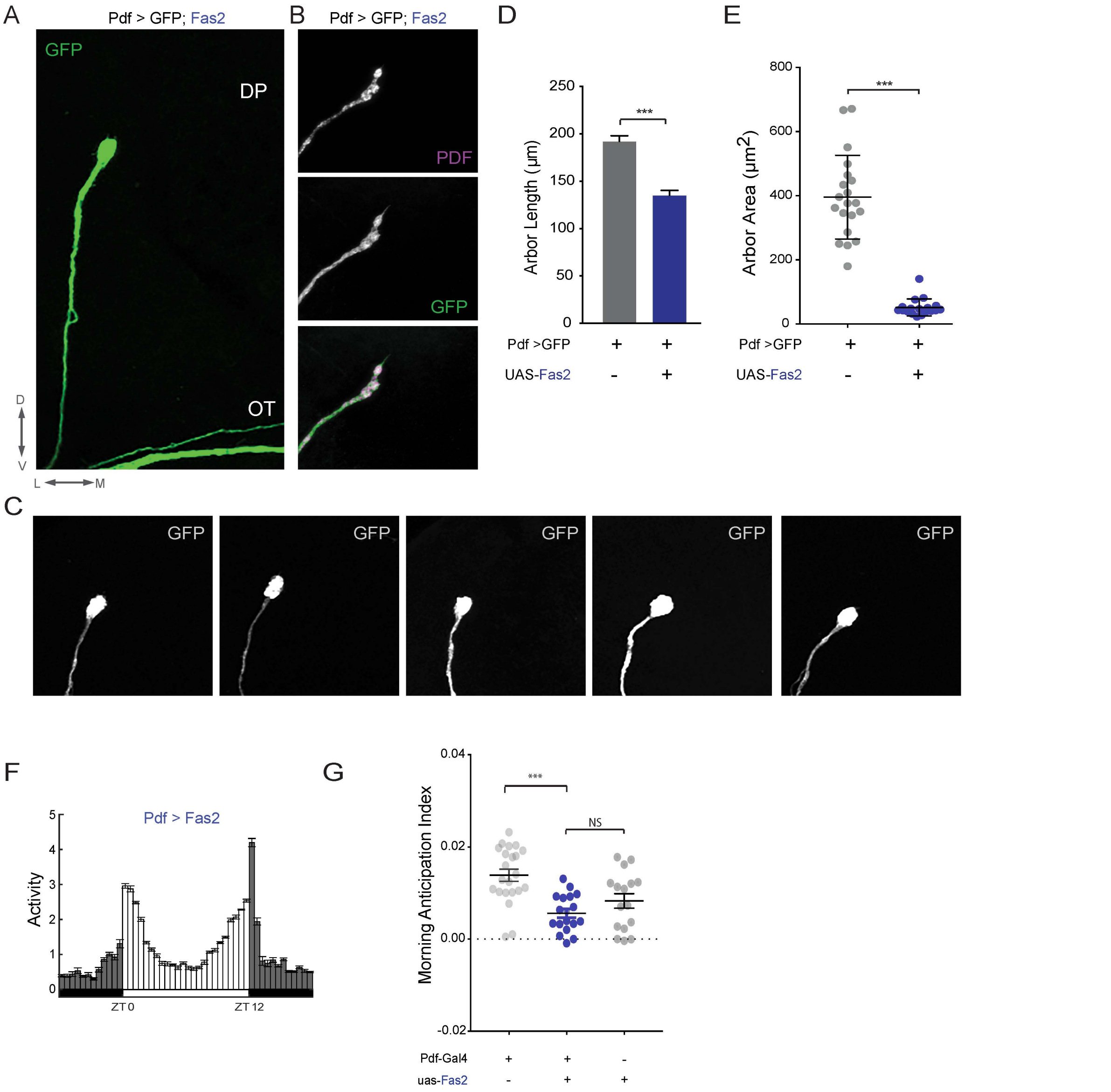

**Figure S5**. Fas2-mediated elimination of the dorsal termini of the s-LNvs does not affect the timing of activity under LD cycles.

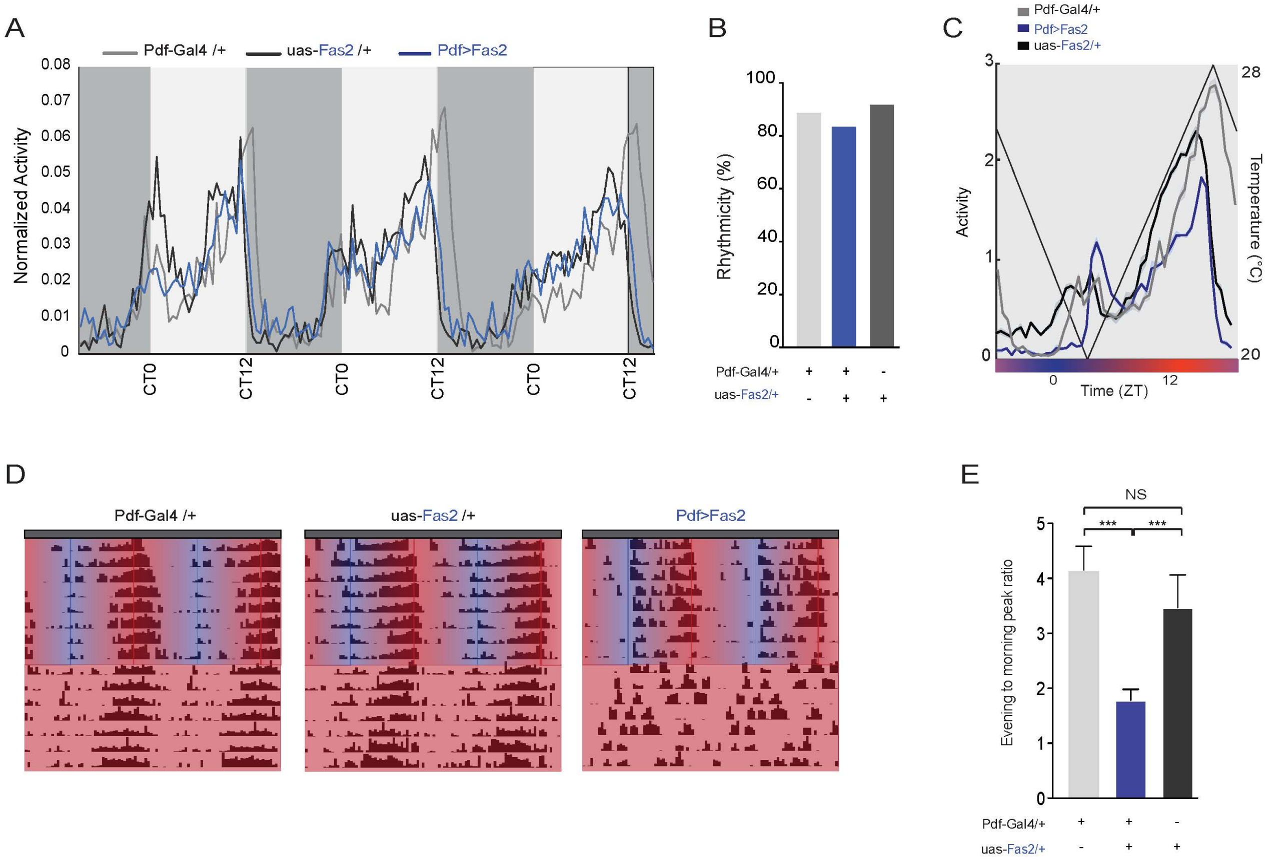

**Figure S6**. Fas2-mediated elimination of the dorsal termini of the s-LNvs does not affect endogenous circadian timekeeping but impairs entrainment to temperature ramps.

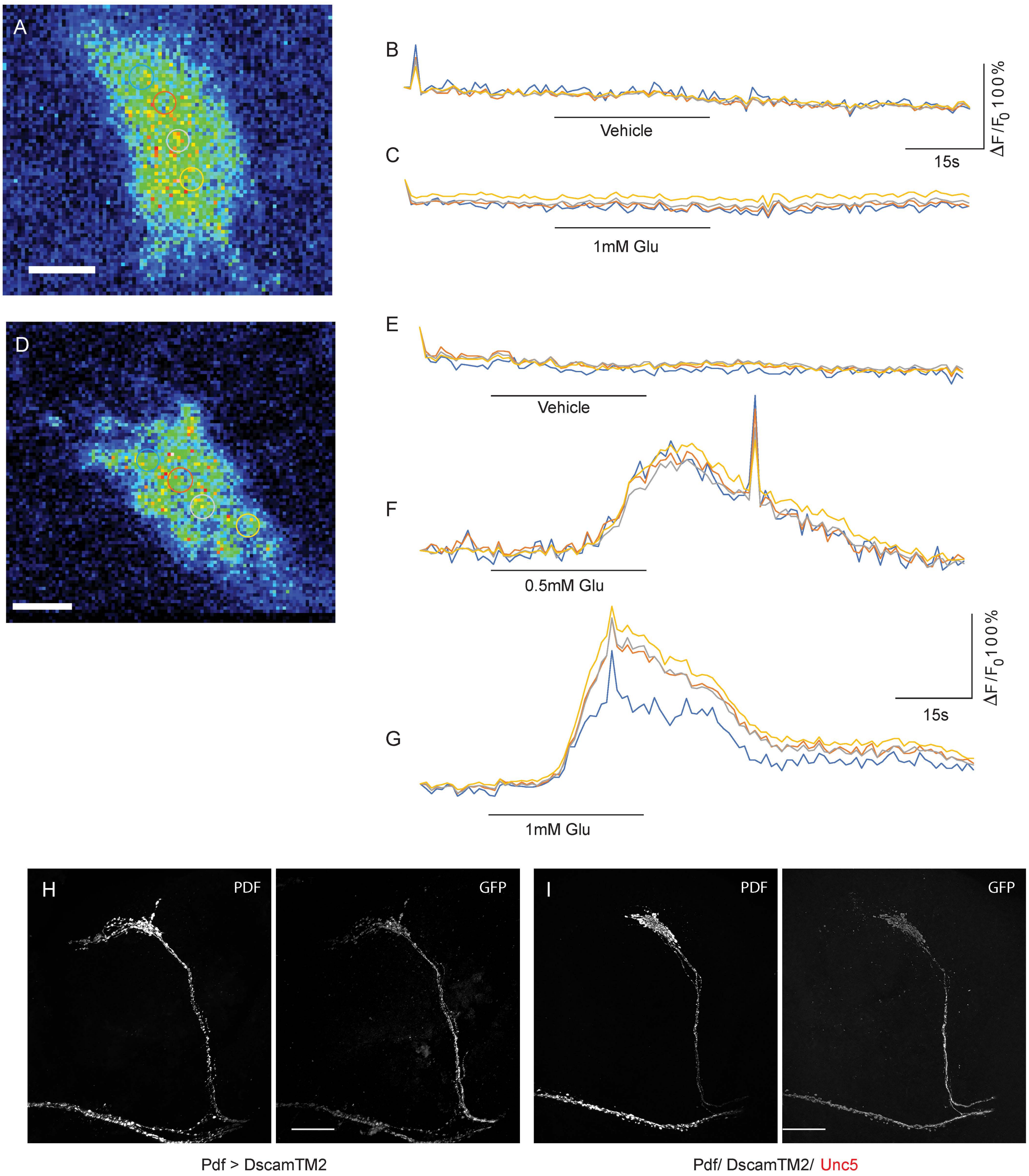

**Figure S7**. Truncated Unc5 expressing s-LNv termini are receptive to glutamate and display rebound excitation following glutamate perfusion.

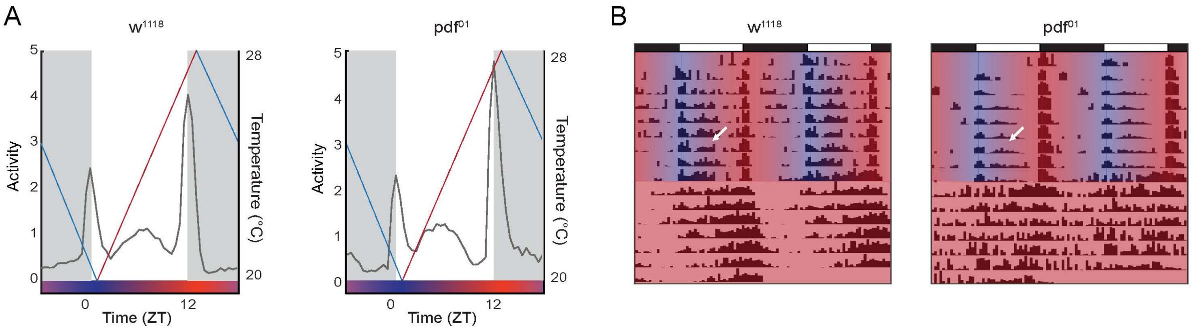

**Figure S8**. The production of a daily, temperature induced mid-day peak under LD cycles does not require PDF.

Table S1.

| Fig 1 | Treatment | Genotype | n | Mean | Std. Deviation | S.E.M. | Passed normality test? | Test | Significant? | Summary | P value |
|---|---|---|---|---|---|---|---|---|---|---|---|
| 1E | Full projection length | | | | | | | unpaired t- test | Significant? | Summary | P value |
| | | Pdf>mCD8:GFP | 22 | 192.10 | 27.45 | 5.85 | yes | Pdf>mCD8:GFP vs. | yes | *** | <0.0001 |
| | | Pdf>Unc5;mCD8:GFP | 16 | 150.20 | 18.89 | 4.72 | yes | Pdf>Unc5;mCD8:GFP | | | |
| 1F | Arbor Area | | | | | | | unpaired t- test | Significant? | Summary | P value |
| | | Pdf>mCD8:GFP | 20 | 395.40 | 130.70 | 29.22 | yes | Pdf>mCD8:GFP vs. | yes | *** | <0.0001 |
| | | Pdf>Unc5;mCD8:GFP | 17 | 95.63 | 23.49 | 5.70 | yes | Pdf>Unc5;mCD8:GFP | | | |
| 1G | x-spread | | | | | | | unpaired t- test | Significant? | Summary | P value |
| | | Pdf>mCD8:GFP | 17 | 52.82 | 10.68 | 2.59 | yes | Pdf>mCD8:GFP vs. | yes | *** | <0.0001 |
| | | Pdf>Unc5;mCD8:GFP | 8 | 17.75 | 5.18 | 1.83 | yes | Pdf>Unc5;mCD8:GFP | | | |
| 1H | y-spread | | | | | | | unpaired t- test | Significant? | Summary | P value |
| | | Pdf>mCD8:GFP | 17 | 25.29 | 5.83 | 1.41 | yes | Pdf>mCD8:GFP vs. | yes | ** | 0.0038 |
| | | Pdf>Unc5;mCD8:GFP | 8 | 17.50 | 5.21 | 1.84 | yes | Pdf>Unc5;mCD8:GFP | | | |
| 1I | z-spread | | | | | | | Mann-Whitney | Significant? | Summary | P value |
| | | Pdf>mCD8:GFP | 17 | 15.88 | 7.16 | 1.74 | no | Pdf>mCD8:GFP vs. | yes | *** | <0.0001 |
| | | Pdf>Unc5;mCD8:GFP | 8 | 7.00 | 1.07 | 0.38 | no | Pdf>Unc5;mCD8:GFP | | | |
| 1J | 3D spread | | | | | | | Mann-Whitney | Significant? | Summary | P value |
| | | Pdf>mCD8:GFP | 17 | 21984 | 12381 | 3003 | yes | Pdf>mCD8:GFP vs. | yes | *** | <0.0001 |
| | | Pdf>Unc5;mCD8:GFP | 8 | 2199 | 1224 | 432.6 | no | Pdf>Unc5;mCD8:GFP | | | |

| Fig 2 | Treatment | Genotype | n | Mean | Std. Deviation | S.E.M. | Passed normality test? | Test | Significant? | Summary | P value | |
|---|---|---|---|---|---|---|---|---|---|---|---|---|
| 2B | Morning Anticipation | | | | | | | Mann Whitney | Significant? | Summary | P value | |
| | | w1118 | 72 | 0.0106 | 0.0058 | 0.0007 | yes | w1118 vs. Pdf 01 | yes | *** | <0.0001 | |
| | | pdf01 | 75 | -1.765E-05 | 0.0056 | 0.0006 | no | | | | | |
| 2D | Morning Anticipation | | | | | | | Kruskal Wallis with Dunn's post test | Significant? | Summary | P Value | |
| | | Pdf-Gal4/+ | 79 | 0.0152 | 0.0080 | 0.0009 | no | Pdf-Gal4/+ vs. Pdf>Unc5 | yes | **** | <0.0001 | A-B |
| | | Pdf>Unc5 | 75 | 0.0062 | 0.0079 | 0.0009 | yes | Pdf-Gal4/+ vs. uas-Unc5/+ | yes | **** | <0.0001 | A-C |
| | | uas-Unc5/+ | 92 | 0.0058 | 0.0059 | 0.0006 | no | Pdf>Unc5 vs. uas-Unc5/+ | no | ns | >0.9999 | B-C |
| 2E | Evening Phase | | | | | | | Kruskal Wallis with Dunn's post test | Significant? | Summary | P Value | |
| | | Pdf-Gal4/+ | 49 | -0.4286 | 0.7706 | 0.1101 | no | Pdf-Gal4/+ vs. Pdf>Unc5 | no | ns | >0.9999 | A-B |
| | | Pdf>Unc5 | 42 | -0.4286 | 0.7034 | 0.1085 | yes | Pdf-Gal4/+ vs. uas-Unc5/+ | yes | **** | <0.0001 | A-C |
| | | uas-Unc5/+ | 48 | -1.365 | 0.8736 | 0.1261 | no | Pdf>Unc5 vs. uas-Unc5/+ | yes | **** | <0.0001 | B-C |

| Fig 3 | Treatment | Genotype | n | Mean | Std. Deviation | S.E.M. | Passed normality test? | Test | Significant? | Summary | P Value | |
|---|---|---|---|---|---|---|---|---|---|---|---|---|
| 3C | Free- Running Period | | | | | | | Kruskall-Wallis + Dunn's post-test | Significant? | Summary | P Value | |
| | | Pdf-Gal4/+ | 55 | 24.0300 | 0.1146 | 0.0155 | no | Pdf-Gal4/+ vs. uas-Unc5/+ | yes | **** | <0.0001 | A-B |
| | | uas-Unc5/+ | 61 | 23.5900 | 0.2142 | 0.0274 | no | Pdf-Gal4/+ vs. uas-DbtL/+ | no | ns | >0.9999 | A-C |
| | | uas-DbtL/+ | 58 | 23.9100 | 0.1977 | 0.0260 | no | Pdf-Gal4/+ vs. Unc5/+;DbtL/+ | yes | **** | <0.0001 | A-D |
| | | Unc5/+;DbtL/+ | 54 | 23.5900 | 0.2394 | 0.0326 | yes | Pdf-Gal4/+ vs. Pdf>Unc5 | no | ns | >0.9999 | A-E |
| | | Pdf>Unc5 | 40 | 24.0400 | 0.2628 | 0.0416 | no | Pdf-Gal4/+ vs. Pdf>DbtL | yes | **** | <0.0001 | A-F |
| | | Pdf>DbtL | 28 | 27.2100 | 0.5998 | 0.1134 | no | Pdf-Gal4/+ vs. Pdf>DbtL;Unc5 | yes | *** | 0.0009 | A-G |
| | | Pdf>DbtL;Unc5 | 58 | 26.2400 | 1.5220 | 0.1999 | no | uas-Unc5/+ vs. uas-DbtL/+ | yes | **** | <0.0001 | B-C |
| | | | | | | | | uas-Unc5/+ vs. Unc5/+;DbtL/+ | no | ns | >0.9999 | B-D |
| | | Kruskall-Wallis + Dunn's post-test | | Significant? | Summary | P Value | | uas-DbtL/+ vs. Pdf>DbtL;Unc5 | yes | **** | <0.0001 | C-G |
| | | uas-Unc5/+ vs. Pdf>Unc5 | | yes | **** | <0.0001 | B-E | Unc5/+;DbtL/+ vs. Pdf>Unc5 | yes | **** | <0.0001 | D-E |
| | | uas-Unc5/+ vs. Pdf>DbtL | | yes | **** | <0.0001 | B-F | Unc5/+;DbtL/+ vs. Pdf>DbtL | yes | **** | <0.0001 | D-F |
| | | uas-Unc5/+ vs. Pdf>DbtL;Unc5 | | yes | **** | <0.0001 | B-G | Unc5/+;DbtL/+ vs. Pdf>DbtL;Unc5 | yes | **** | <0.0001 | D-G |
| | | uas-DbtL/+ vs. Unc5/+;DbtL/+ | | yes | *** | 0.0001 | C-D | Pdf>Unc5 vs. Pdf>DbtL | yes | **** | <0.0001 | E-F |
| | | uas-DbtL/+ vs. Pdf>Unc5 | | no | ns | >0.9999 | C-E | Pdf>Unc5 vs. Pdf>DbtL;Unc5 | yes | ** | 0.0031 | E-G |
| | | uas-DbtL/+ vs. Pdf>DbtL | | yes | **** | <0.0001 | C-F | Pdf>DbtL vs. Pdf>DbtL;Unc5 | no | ns | 0.4648 | F-G |

| Fig 3 | Treatment | Genotype | n | Mean | Std. Deviation | S.E.M. | Passed normality test? | Test | | Watson-Wheeler | Watson's Two-Sample |
|---|---|---|---|---|---|---|---|---|---|---|---|
| 3D | Circular Phase Analysis | | | | | | | Both circular statistics tests | Significant? | P-value | P-value |
| | | Pdf-Gal4 /+; | 53 | 0.0377 | 0.7522 | 0.1033 | N/A | Pdf>Unc5 vs. Pdf-Gal4/+ | no | 0.167 | <0.10 |
| | | uas-Unc5/+ | 60 | -1.4250 | 0.6299 | 0.0813 | N/A | Pdf>Unc5 vs. uasUnc5 | yes | <0.0001 | <0.001 |
| | | Pdf-Gal4 /+;Unc5/+ | 41 | 0.4268 | 0.9392 | 0.1467 | N/A | Pdf>DbtL vs. Pdf-Gal4 | yes | <0.0001 | <0.001 |
| | | uas-DbtL/+ | 59 | -0.9068 | 0.8172 | 0.1064 | N/A | Pdf>DbtL vs. uas-DbtL | yes | <0.0001 | <0.001 |
| | | uas-DbtL/+;uas-Unc5/+ | 49 | -1.3270 | 0.9385 | 0.1341 | N/A | Pdf>Unc5; DbtL vs. Pdf>Unc5 | yes | 0.0002905 | <0.001 |
| | | Pdf-Gal4 /uas-DbtL | 35 | 2.0140 | 0.8444 | 0.1427 | N/A | Pdf>Unc5; DbtL vs. Pdf>DbtL | no | 0.4324 | 0.113 |
| | | Pdf-Gal4 /DbtL;Unc5/+ | 60 | 1.3000 | 1.1760 | 0.1518 | N/A | Pdf>Unc5; DbtL vs. Pdf-Gal4 | yes | <0.0001 | <0.001 |
| | | | | | | | | Pdf>Unc5; DbtL vs. uas-DbtL | yes | <0.0001 | <0.001 |
| | | | | | | | | Pdf>Unc5; DbtL vs. uasUnc5 | yes | <0.0001 | <0.001 |

| Fig 5 | Treatment | Genotype | n | Mean | Std. Deviation | S.E.M. | Passed normality test? | Test | | | | |
|---|---|---|---|---|---|---|---|---|---|---|---|---|
| 5B | Evening/morning ratio | | | | | | | Kruskal Wallis with Dunn's post test | Significant? | Summary | P Value | |
| | | uas-Unc5/+ | 31 | 2.5490 | 1.1250 | 0.2020 | no | Unc5/+ vs. Pdf-Gal4/+ | no | ns | 0.0749 | A-C |
| | | Pdf>Unc5 | 32 | 1.3470 | 0.4999 | 0.0884 | yes | Unc5/+ vs. Unc5/Pdf-Gal4 | yes | **** | <0.0001 | B-C |
| | | Pdf-Gal4/+ | 32 | 4.2550 | 2.4420 | 0.4327 | yes | Pdf-Gal4/+ vs. Unc5/Pdf-Gal4 | yes | **** | <0.0001 | A-B |
| 5D | Heating Index | | | | | | | Kruskal Wallis with Dunn's post test | Significant? | Summary | P Value | |
| | | uas-Unc5/+ | 59 | 0.4729 | 0.3111 | 0.0405 | yes | Pdf-Gal4/+ vs. Pdf>Unc5 | yes | *** | <0.0001 | B-C |
| | | Pdf>Unc5 | 58 | 0.1093 | 0.3336 | 0.0503 | no | Pdf-Gal4/+ vs. uas-Unc5/+ | no | NS | 0.4136 | A-B |
| | | Pdf-Gal4/+ | 44 | 0.5667 | 0.2393 | 0.0314 | yes | Pdf>Unc5 vs. uas-Unc5/+ | yes | *** | <0.0001 | A-C |
| 5F | Heating Index | | | | | | | Kruskal Wallis with Dunn's post test | Significant? | Summary | P Value | |
| | | Pdf-Gal4/+ | 58 | 0.5667 | 0.2393 | 0.0314 | no | Pdf-Gal4/+ vs. Pdf>uas-hid | yes | *** | 0.0002 | A-B |
| | | Pdf>uas-hid | 12 | -0.1826 | 0.4685 | 0.1353 | yes | Pdf-Gal4/+ vs. uas-hid/+ | yes | * | 0.0151 | A-C |
| | | uas-hid/+ | 16 | 0.7542 | 0.1718 | 0.0429 | no | Pdf>uas-hid vs. uas-hid/+ | yes | **** | <0.0001 | B-C |
| 5H | Heating Index | | | | | | | Mann Whitney | | Summary | P Value | |
| | | w1118 | 57 | 0.5951 | 0.2188 | 0.0290 | no | w118 vs pdf01 | no | NS | 0.2981 | |
| | | pdf01 | 55 | 0.5051 | 0.3448 | 0.0465 | no | | | | | |

| Fig 6 | Treatment | Genotype | n | Mean | Std. Deviation | S.E.M. | Passed normality test? | Test | | | | |
|---|---|---|---|---|---|---|---|---|---|---|---|---|
| 6I | Heating Index | | | | | | | Kruskal Wallis with Dunn's post test | Significant? | Summary | P Value | |
| | | Pdf-Gal4/+ | 30 | 0.3845 | 0.2504 | 0.0457 | yes | Pdf-Gal4 /+vs. Pdf>GluClRNAi | yes | ** | 0.0042 | A-B |
| | | Pdf>dicer2; GluCl RNAi | 23 | -0.0391 | 0.349 | 0.0727 | yes | Pdf-Gal4 /+vs. uas-GluCl RNAi/+ | yes | *** | 0.0002 | A-C |
| | | dicer2/+;uas-GluCl RNAi/+ | 31 | 0.679 | 0.1843 | 0.0331 | no | Pdf>GluClRNAi vs. uas-GluClRNAi/+ | yes | *** | <0.0001 | B-C |

| Fig 7 | Treatment | Genotype | n | Mean | Std. Deviation | S.E.M. | Passed normality test? | Test | | | | |
|---|---|---|---|---|---|---|---|---|---|---|---|---|
| 7B | Mid-day Activity | | | | | | | Kruskal Wallis with Dunn's post test | Significant? | Summary | P Value | |
| | | uas-Unc5/+ | 62 | 0.7140 | 0.4524 | 0.0575 | yes | Unc5/+ vs. Pdf-Gal4/+ | yes | * | 0.0343 | A-B |
| | | Pdf>Unc5 | 62 | 0.3082 | 0.1927 | 0.0245 | no | Unc5/+ vs. Pdf-Gal4/Unc5 | yes | **** | <0.0001 | A-C |
| | | Pdf-Gal4/+ | 61 | 1.0750 | 0.7129 | 0.0913 | no | Pdf-Gal4/+ vs. Pdf-Gal4/Unc5 | yes | **** | <0.0001 | B-C |
| 7D | Mid-day Activity | | | | | | | Mann Whitney | | Summary | P Value | |
| | | w1118 | 60 | 0.9984 | 0.4630 | 0.0598 | no | w1118 vs. pdf01 | | NS | 0.6302 | |
| | | pdf01 | 58 | 0.9599 | 0.4846 | 0.0636 | no | | | | | |

| Fig S1 | Treatment | Genotype | n | Mean | Std. Deviation | S.E.M. | Passed normality test? | Test | | | |
|---|---|---|---|---|---|---|---|---|---|---|---|
| S1C | Max. # Neurites | | | | | | | unpaired t-test | Significant? | Summary | P value |
| | | Pdf>mCD8:GFP | 12 | 2.2500 | 0.4523 | 0.1306 | yes | Pdf>mCD8:GFP vs. | yes | *** | 0.0005 |
| | | Pdf>Unc5;mCD8:GFP | 8 | 3.3750 | 0.7440 | 0.2631 | yes | Pdf>Unc5;mCD8:GFP | | | |

| Fig S3 | Treatment | Genotype | n | Mean | Std. Deviation | S.E.M. | Passed normality test? | Test | | | | |
|---|---|---|---|---|---|---|---|---|---|---|---|---|
| S3A | Morning Phase | | | | | | | AnoVA + Tukey's post test | | | P Value | |
| | | Pdf-Gal4/+ | 49 | -0.02041 | 1.407 | 0.201 | yes | PdfGal4/+ vs. Pdf>Unc5 | yes | * | 0.0265 | A-B |
| | | Pdf>Unc5 | 42 | 0.7024 | 1.339 | 0.2066 | yes | PdfGal4/+ vs. uas-Unc5/+ | yes | ** | 0.0078 | A-C |
| | | uas-Unc5/+ | 48 | 0.7917 | 1.184 | 0.1709 | yes | Pdf>Unc5 vs. uas-Unc5/+ | no | ns | 0.9445 | B-C |
| S3C | Evening Anticipation | | | | | | | Unpaired t-test | | Summary | P Value | |
| | | w1118 | 72 | 0.0165 | 0.0102 | 0.0012 | yes | w1118 vs. Pdf 01 | | ns | 0.0767 | |
| | | pdf01 | 75 | 0.0195 | 0.0103 | 0.0012 | yes | | | | | |
| S3D | Evening Anticipation | | | | | | | Kruskal Wallis with Dunn's | Significant? | Summary | P Value | |
| | | Pdf-Gal4/+ | 79 | 0.0112 | 0.0147 | 0.0016 | yes | Pdf-Gal4/+ vs. Pdf>Unc5 | no | ns | 0.1635 | A-B |
| | | Pdf>Unc5 | 75 | 0.0164 | 0.0132 | 0.0015 | no | Pdf-Gal4/+ vs. uas-Unc5/+ | yes | **** | <0.0001 | A-C |
| | | uas-Unc5/+ | 92 | 0.0263 | 0.0121 | 0.0013 | yes | Pdf>Unc5 vs. uas-Unc5/+ | yes | **** | <0.0001 | B-C |

| Fig S5 | Treatment | Genotype | n | Mean | Std. Deviation | S.E.M. | Passed normality test? | Test | | | | |
|---|---|---|---|---|---|---|---|---|---|---|---|---|
| S5D | Arbor Length | | | | | | | Unpaired t-test | Significant? | Summary | P Value | |
| | | Pdf>mCD8:GFP | 22 | 192.1 | 27.5 | 5.9 | yes | Pdf>mCD8:GFP vs | yes | *** | <0.0001 | |
| | | Pdf>Fas2;mCD8:GFP | 16 | 135.2 | 20.5 | 5.1 | yes | Pdf>Fas2; mCD8:GFP | | | | |
| S5E | Arbor Area | | | | | | | Mann-Whitney test | Significant? | Summary | P Value | |
| | | Pdf>mCD8:GFP | 20 | 395.5 | 130.7 | 29.2 | yes | Pdf>mCD8:GFP vs | yes | *** | <0.0001 | |
| | | Pdf>Fas2;mCD8:GFP | 18 | 51.4 | 26.6 | 6.3 | no | Pdf>Fas2; mCD8:GFP | | | | |
| S5G | Morning Anticipation | | | | | | | AOVAVA with Tukey's post-hoc test | Significant? | Summary | P Value | |
| | | Pdf-Gal4/+ | 22 | 0.0139 | 0.0061 | 0.0013 | yes | uas-Fas2/+ vs. Pdf>Fas2 | no | ns | 0.348 | A-B |
| | | uas-Fas2/+ | 18 | 0.0056 | 0.0040 | 0.0010 | yes | uas-Fas2/+ vs. Pdf-Gal4/+ | yes | * | 0.0103 | A-C |
| | | Pdf>Fas2 | 16 | 0.0083 | 0.0063 | 0.0016 | yes | Pdf>Fas2 vs. Pdf-Gal4/+ | yes | **** | <0.0001 | B-C |

| Fig S6 | Treatment | Genotype | n | Mean | Std. Deviation | S.E.M. | Passed normality test? | Test | | | | |
|---|---|---|---|---|---|---|---|---|---|---|---|---|
| S6E | Evening/morning ratio | | | | | | | Kruskal Wallis with Dunn's | Significant? | Summary | P Value | |
| | | Pdf-Gal4/+ | 32 | 4.1550 | 2.4420 | 0.4317 | yes | uas-Fas2/+ vs. Pdf-Gal4/+ | no | ns | >0.9999 | A-B |
| | | Pdf>Fas2 | 32 | 1.7380 | 0.5315 | 0.0940 | yes | uas-Fas2/+ vs. Pdf>Fas2 | yes | **** | <.0001 | A-C |
| | | uas-Fas2/+ | 31 | 4.0600 | 1.9000 | 0.3413 | no | Pdf-Gal4/+ vs. Pdf>Fas2 | yes | **** | <.0001 | B-C |
